# Molecular principles for graded activation of dopamine D1 receptor

**DOI:** 10.64898/2026.07.28.741251

**Authors:** Xinyue Zhang, Yu Zheng, Heng Liu, Jingjing Hou, Luyu Fan, Xinheng He, Jun Sun, Tao Liu, Jianhui Zhou, Ruxin Lei, Mingyang Li, Wen Hu, Xi Cheng, Sheng Wang, Xin Xie, H. Eric Xu, Shimeng Guo, Youwen Zhuang

**Affiliations:** State Key Laboratory of Drug Research, Shanghai Institute of Materia Medica, Chinese Academy of Sciences, Shanghai 201203, China; School of Chinese Materia Medica, Nanjing University of Chinese Medicine, Nanjing 210046, China; School of Pharmaceutical Science and Technology, Hangzhou Institute for Advanced Study, University of Chinese Academy of Sciences, Hangzhou 310024, China; University of Chinese Academy of Sciences, Beijing 100049, China; Key Laboratory of Systems Health Science of Zhejiang Province, School of Life Science, Hangzhou Institute for Advanced Study, University of Chinese Academy of Sciences, Hangzhou 310024, China; School of Life Sciences and Technology, Tongji University, Shanghai 200092, China; Department of Pharmaceutical and Artificial-Intelligence Sciences, School of Medicine, Shanghai Jiao Tong University, Shanghai 200025, China; Key Laboratory of Cell Differentiation and Apoptosis of Chinese Ministry of Education, School of Medicine, Shanghai Jiao Tong University, Shanghai 200025, China; Key Laboratory of Multi-Cell Systems, Shanghai Institute of Biochemistry and Cell Biology, Center for Excellence in Molecular Cell Science, Chinese Academy of Sciences, University of Chinese Academy of Sciences, Shanghai 200031, China; Artificial Intelligence Clinical Research Center for drug discovery, Shanghai Key Laboratory of Flexible Medical Robotics, Tongren Hospital, School of Medicine, Shanghai Jiao Tong University, Shanghai 200025, China

## Abstract

G protein-coupled receptors (GPCRs) signal across a continuum of activation states, yet how ligands encode distinct signaling efficacies remains poorly understood. Here, we define the molecular mechanism underlying graded activation of the dopamine D1 receptor (D1R), a major therapeutic target for neuropsychiatric disorders. Functional analyses reveal that two widely used pharmacological tools, LE300 and SCH23390, possess intrinsic efficacy as an inverse agonist and a weak partial agonist, respectively, rather than the efficacy-silent neutral antagonists. Structural, molecular dynamics and mutagenesis analyses capture previously unrecognized inactive and intermediate receptor activation states that bridge known active conformations, and reveal that ligand efficacy is encoded through the progressive engagement of a conserved activation pathway centered on the W^6.48^ toggle switch. Guided by this mechanism, a single chemical modification markedly increases the agonist efficacy of SCH23390. Comparison with dopamine D2 receptor structures further reveals a conserved mechanism of inverse agonism despite distinct subtype-specific recognition. Together, these findings establish a structural framework for graded agonism at D1R and provide general principles for the rational design of efficacy-tuned therapeutics at dopamine receptors and related GPCRs.

## Introduction

Dopamine signaling is a critical component of the central and peripheral nervous systems, mediating essential physiological processes including motor control, reward processing, cognitive function, and blood pressure regulation. The signaling of dopamine is orchestrated through five distinct dopamine receptor subtypes (D1R-D5R), which belong to the G protein-coupled receptor (GPCR) family and are classified into two families, the D1-like receptors (D1R and D5R) and D2-like receptors (D2R, D3R, and D4R). Among them, D1R is the most abundantly expressed dopamine receptor in the CNS^1,2^. D1R primarily signals through stimulatory G protein G_s/olf_ to activate adenylyl cyclase and elevate intracellular cyclic adenosine monophosphate (cAMP) levels. Dysregulation of D1R signaling has been implicated in various neurological and psychiatric disorders, including Parkinson’s disease (PD), schizophrenia, and substance abuse disorders, highlighting its significance as a crucial therapeutic target^3^. Unraveling its signaling mechanisms is essential for understanding both the therapeutic and side effects of dopaminergic drug.

Pharmacological modulation of D1R spans a continuous spectrum of ligand efficacies that correspond to distinct receptor conformational states and signaling outputs. Full agonists such as SKF81297 stabilize fully active conformations and drive robust G_s_ signaling comparable to the endogenous ligand dopamine^4^. Several FDA-approved dopaminergic drugs, including apomorphine and rotigotine, also show full agonist activity at D1R *in vitro*, although their clinical effects are primarily attributed to D2-like receptor activation^4–6^. In contrast, inverse agonists preferentially stabilize inactive receptor states and suppress constitutive D1R activity. Notably, several antipsychotic drugs, including chlorpromazine, fluphenazine, and clozapine, have been reported to exhibit inverse agonism at D1R^7,8^. Neutral antagonists, a category that has long included SCH23390^9–11^ and LE300^12^, occupy the orthosteric binding pocket (OBP) without intrinsic efficacy and competitively block agonist-induced activation. Between these functional extremes, partial agonists such as fenoldopam, SKF83959 and SKF38393, elicit submaximal receptor activation even at saturating concentrations. Rather than fully shifting the receptor population toward a single active state, partial agonists are thought to stabilize intermediate conformational states that support limited G protein engagement and attenuated downstream signaling^13–15^. This constrained efficacy profile may preserve aspects of physiological dopaminergic tone while avoiding the excessive signaling associated with full agonism or the functional silencing imposed by inverse agonism^16–18^. As such, D1R partial agonists offer a pharmacological strategy to fine-tune receptor activation, potentially achieving a balance between therapeutic efficacy, tolerability, and maintenance of signaling homeostasis, as exemplified by the clinical development of tavapadon^16,19,20^.

Despite the clinical relevance of D1R modulation, current molecular understanding of its activation mechanisms remains incomplete. Recent structure pharmacological studies including those from our group have primarily focused on the agonist-bound active states of D1R, elucidating the molecular mechanisms of agonist recognition, positive allosteric modulation, biased signaling, and selective G_s_ coupling at D1R^4,21–26^. Despite these progresses, a comprehensive and precise understanding of D1R’s conformational landscape and the nuanced roles of ligands in modulating its activity is still lacking, mainly attributes to a lack of inactive D1R structure, limiting the rational design of therapeutics with tailored efficacy to modulate receptor signaling and treat diseases requiring precise modulation of dopaminergic tone.

In this study, we sought to define how ligands spanning the full efficacy spectrum modulate D1R activation. We first systematically reassessed the pharmacological properties of several widely used D1R ligands and found that two classical antagonists, LE300 and SCH23390, instead occupy distinct positions within the efficacy spectrum. These ligands provided an opportunity to investigate receptor states that had remained inaccessible to structural and mechanistic analyses. We then determine cryo-electron microscopy (cryo-EM) structures of D1R capturing the inactive state stabilized by LE300 and the intermediate active state induced by SCH23390. Together with molecular dynamics simulations and mutagenesis, these results delineate how progressively different ligand-receptor interactions propagate through a conserved activation pathway to produce graded signaling responses. Finally, structure-guided ligand redesign experimentally validates this mechanism by converting SCH23390 into a high-efficacy partial agonist through a single chemical modification. Together, our findings uncover how ligand efficacy is encoded in D1R, establish a mechanistic framework for graded GPCR activation, and provide principles for the rational design of efficacy-tuned therapeutics.

## Results

### Revisiting D1R pharmacology reveals a spectrum of receptor activation states

D1R pharmacology has traditionally relied on several prototypical antagonists, including SCH23390 and LE300, to define receptor blockade in both cellular and animal models^9,12,27^. Using cAMP signaling analyses, we first reassessed the intrinsic activities of these widely used ligands. In contrast to their canonical antagonist designation, we found that these compounds exhibit distinct intrinsic activities and do not function as neutral antagonists under our experimental conditions. SCH23390 displayed modest but reproducible agonist activity relative to dopamine, consistent with weak partial agonism (EC_50_ = 4.85 nM; Emax = 26.4%), whereas LE300 suppressed basal D1R signaling, exhibiting inverse agonist properties (EC_50_ = 19.8 nM; Emax = −13.8%) (Figs. 1a-c and Supplementary Table 1). In addition, several other antipsychotic agents historically classified as D1R antagonists such as mesoridazine and periciazine also suppressed basal D1R activity, therefore better characterized as inverse agonists rather than neutral antagonists (Extended Data Fig. 1 and Supplementary Table 2). Chlorpromazine, fluphenazine, and haloperidol showed the same pattern, in agreement with prior reports of their inverse agonist activity at D1R^7,8^. These findings redefine the functional identities of established D1R ligands and demonstrate that conventional antagonist designation does not fully capture their intrinsic pharmacological properties.

**Fig. 1.**
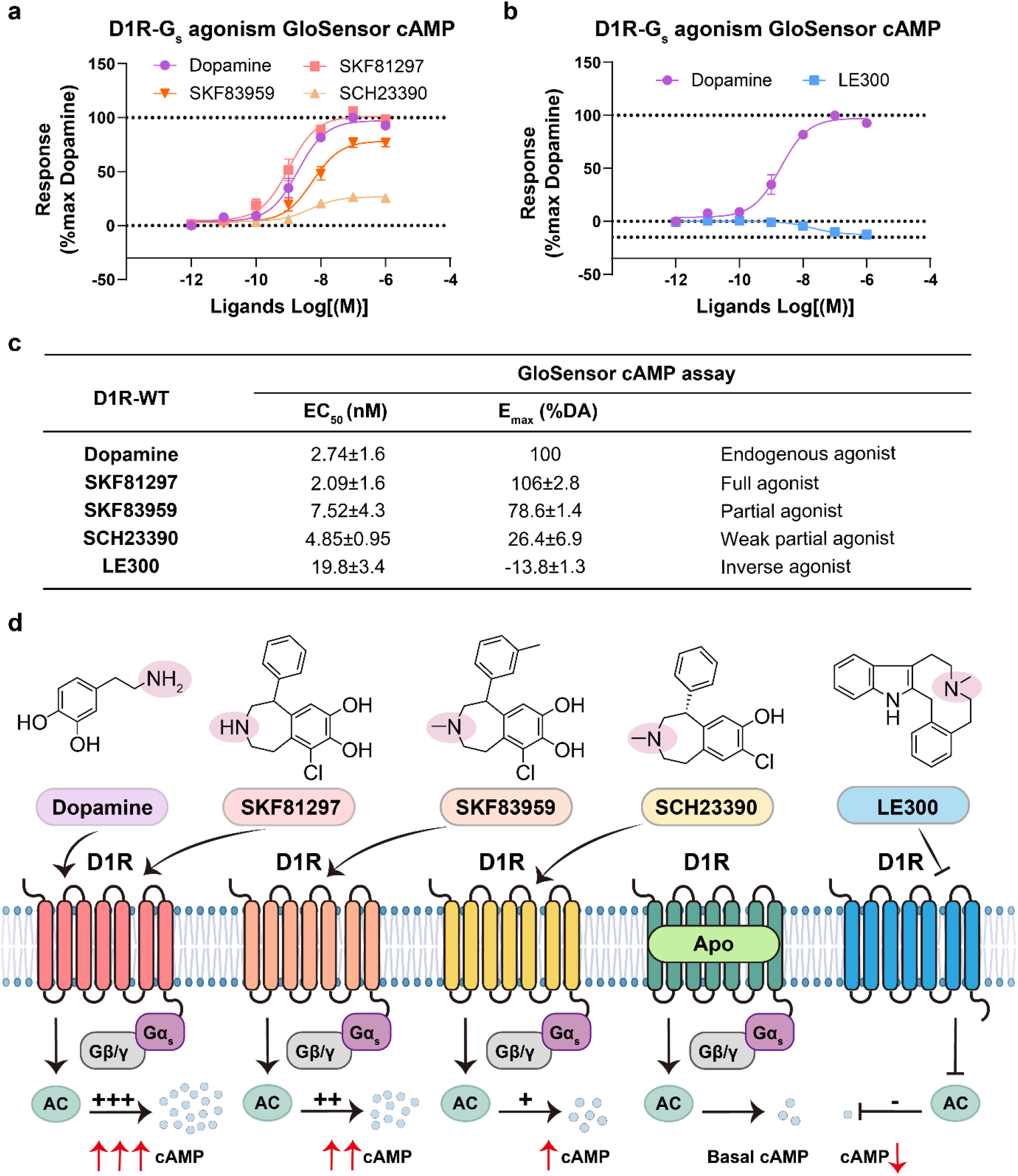
D1R ligands span a continuum from inverse agonism to full agonism. **a,** Concentration-response curves for dopamine, SKF81297, SKF83959 and SCH23390 measured using the GloSensor cAMP assay in HEK293 cells expressing wild-type (WT) D1R. Responses were normalized to the maximal dopamine response (100%), with apo basal activity defined as 0%. Data are mean ± S.E.M. from three independent experiments (*n* = 3), each performed in triplicate. **b,** Concentration-response curves for dopamine and LE300 measured using the GloSensor cAMP assay in HEK293 cells expressing WT D1R. Responses were normalized to the maximal dopamine response (100%), with apo basal activity defined as 0%. Data are mean ± S.E.M. from three independent experiments (*n* = 3), each performed in triplicate. **c,** Summary of EC_50_, Emax and pharmacological activity for the five ligands. Responses were normalized to the maximal dopamine response (100%), with apo basal activity defined as 0%. Data are mean ± S.E.M. from three independent experiments (*n* = 3), each performed in triplicate. DA, dopamine. **d,** Chemical structures and schematic representation of the effects of dopamine, SKF81297, SKF83959, SCH23390 and LE300 on D1R-mediated G_s_ signaling and cAMP production. Apo D1R (no ligand, green) defines basal activity.

Building on this functional reassessment, we next characterized a chemically diverse ligand panel spanning a broad efficacy spectrum, including dopamine, the SKF-series agonists SKF81297 and SKF83959, SCH23390, and LE300 (Fig. 1d). LE300 was selected as a representative inverse agonist because of its high potency and established D1-like receptor selectivity. Despite engaging the OBP, these ligands elicited markedly different D1R signaling responses. Dopamine and SKF81297 produced near-maximal receptor activation, consistent with full agonism (EC_50_ = 2.09 nM, Emax = 106% for SKF81297), whereas SKF83959 exhibited partial agonism (EC_50_ = 7.52 nM, Emax = 78.6%) (Fig. 1a, c and Supplementary Table 1). Together with the redefined activities of SCH23390 and LE300, these ligands establish a functional continuum ranging from inverse agonism to weak partial, partial and full activation, providing a pharmacologically resolved framework for capturing and comparing the distinct D1R conformational states underlying different signaling outputs (Fig. 1d).

### LE300 locks D1R into an inactive conformation

To capture the inactive state of D1R, we engineered a stabilized receptor construct by replacing the flexible intracellular loop 3 (ICL3) with BRIL and introducing additional stabilizing mutations. This strategy enabled us to determine the cryo-EM structure of D1R in complex with the inverse agonist LE300 at an overall resolution of 3.6 Å, providing sufficient detail to define the receptor architecture, ligand-binding pose, and ligand-receptor interactions (Figs. 2a-b, Extended Data Figs. 2f-h and Supplementary Table 3). This structure, to our knowledge, represents the first direct visualization of D1R in an inactive conformation.

**Fig. 2.**
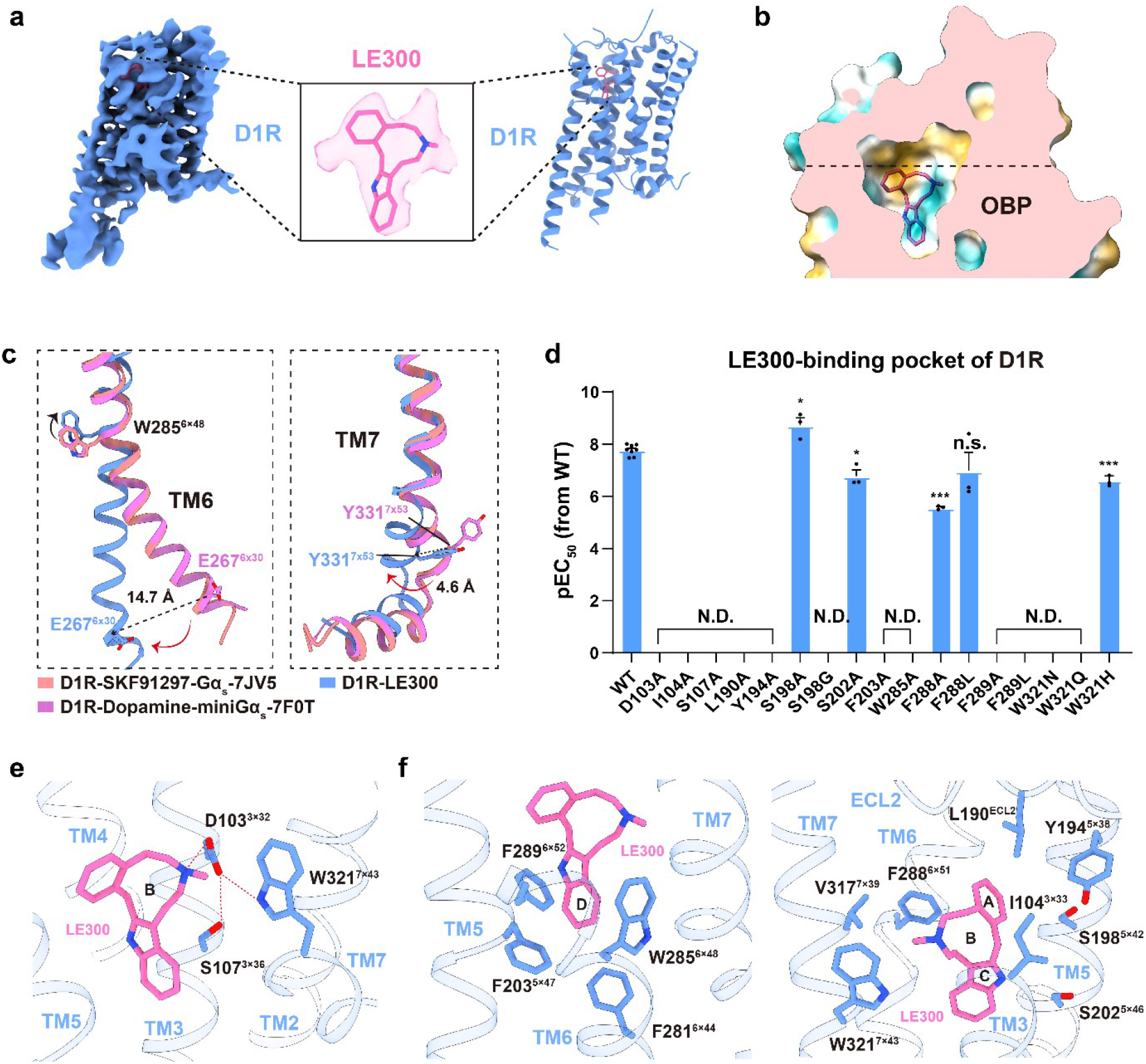
Structural basis of LE300 recognition by inactive D1R. **a,** Cryo-EM density map (left) and cartoon representation (right) of the LE300-D1R complex. The inset shows LE300 and its cryo-EM density. **b,** Cutaway surface representation of the LE300-binding pocket in D1R, colored by hydrophilic (cyan) and hydrophobic (yellow) character. OBP, orthosteric binding pocket. **c,** Close-up views of the intracellular ends of TM6 and TM7. The Cα atom of E267^6.30^ shifts by 14.7 Å and that of Y331^7.53^ shifts by 4.6 Å between the LE300-bound inactive state and the active structures. **d,** Potencies of LE300 for suppressing constitutive cAMP signaling at WT D1R and the indicated binding-pocket mutants, measured using the HTRF cAMP assay in HEK293 cells. Each point represents an independent experiment; data are mean ± S.E.M. from three to nine independent experiments (WT, *n* = 9; mutants, *n* = 3), each performed in triplicate. N.D., not determined, denotes either no response or a response for which reliable parameter estimates could not be established over the tested concentration range. For variants with quantifiable responses, *P* values were determined by one-way ANOVA with Dunnett’s multiple-comparisons test versus WT. \**P* < 0.05, \*\**P* < 0.01, \*\*\**P* < 0.001 and \*\*\*\**P* < 0.0001; n.s., not significant. **e-f,** Views of the LE300-binding pocket. The polar network formed by D103^3.32^, S107^3.36^ and W321^7.43^ is shown in (**e**); two views of hydrophobic contacts surrounding the indole and distal aromatic portions of LE300 are shown in (**f**). Red dashed lines indicate polar interactions.

The LE300-bound D1R structure adopts a compact receptor architecture characteristic of inactive GPCR states, featuring a closed intracellular cavity that is incompatible with G protein coupling (Extended Data Fig. 3a). Structural comparison with active D1R structures bound to the full agonists SKF81297 and dopamine extensive rearrangements across both extracellular and intracellular regions of the receptor^4,26^. Notably, the extracellular loop 2 (ECL2) bends toward the transmembrane domain (TMD) core in the inactive state, forming a lid-like structure over the OBP (Fig.2b and Extended Data Fig. 3b). At the intracellular side, TM6 rotates around the conserved toggle switch residue W285^6×48^ and undergoes a pronounced inward displacement, with the Cα atom of E267^6×30^ shifting by 14.7 Å relative to the active state (Fig. 2c). Conversely, the intracellular end of TM7 moves outward, with a 4.6 Å displacement measured at the Cα atom of Y331^7×53^ (Fig. 2c). Additional rearrangements, including inward shifts of the extracellular regions of TM1, TM2, and TM3, further reshape the receptor core (Extended Data Fig. 3c). These coordinated rearrangements define the structural features associated with the inactive D1R state.

LE300 is deeply buried within the OBP of D1R, engaging extensive interactions with residues contributed by multiple transmembrane helices (TMs) 3, 5, 6, and 7 (Figs. 2b, e-f, and Extended Data Fig. 3b). The ligand adopts an elongated conformation and is anchored by a conserved ionic interaction between its protonated amine and D103^3×32^, a hallmark interaction shared by aminergic GPCR ligands (Fig. 2e). This interaction is further reinforced by a polar network involving S107^3×36^ and W321^7×43^ (Fig. 2e), which effectively stabilizes the central scaffold of LE300. The indole ring of LE300 inserts deeply into a hydrophobic chamber near the TMD center, forming extensive packing interactions with surrounding residues such as F203^5×47^, F281^6×44^, W285^6×48^, and F289^6×52^ (Fig. 2f). Additional hydrophobic residues, including I104^3×33^, L190^ECL2^, F288^6×51^, and W321^7×43^, further shape the ligand-binding environment and accommodate the distal aromatic moieties of the ligand (Fig. 2f). Consistent with this structural framework, mutations of residues involved in ligand anchoring and pocket formation, including D103^3×32^ and W321^7×43^, as well as hydrophobic residues surrounding W285^6×48^, substantially impair LE300 binding and function (Fig. 2d, Extended Data Fig. 4a and Supplementary Table 4). These results indicate that polar anchoring and hydrophobic packing act cooperatively to stabilize the LE300-bound inactive state of D1R (Extended Data Fig. 3c).

### Recognition of SCH23390 in D1R

To elucidate how SCH23390 produces weak partial agonist activity, we determined the cryo-EM structure of the SCH23390-bound D1R-G_s_ complex at an overall resolution of 3.2 Å (Fig. 3a, Extended Data Figs. 2n-p and Supplementary Table 3). The receptor adopts an active conformation closely resembling previously reported agonist-bound D1R-G_s_ structures, with Cα RMSD values of 0.38-0.86 Å following receptor alignment. The well-resolved cryo-EM density enabled unambiguous modeling of SCH23390 within the OBP (Fig. 3a and Extended Data Fig. 2o). In the structure, SCH23390 is fully buried into the OBP buried in the OBP and adopts an L-shaped configuration that closely overlaps those of the benzazepine agonists SKF81297, SKF83959, and fenoldopam^4,25^ (Extended Data Fig. 5a). Its benzazepine scaffold is sandwiched between TM3 and TM6, with the protonated amine forming the conserved ionic interaction with D103^3×32^ (Fig. 3c). As observed for other D1R ligands, this interaction is reinforced by a polar network involving S107^3×36^ and W321^7×43^, anchoring SCH23390 within the OBP (Fig. 3c).

**Fig. 3.**
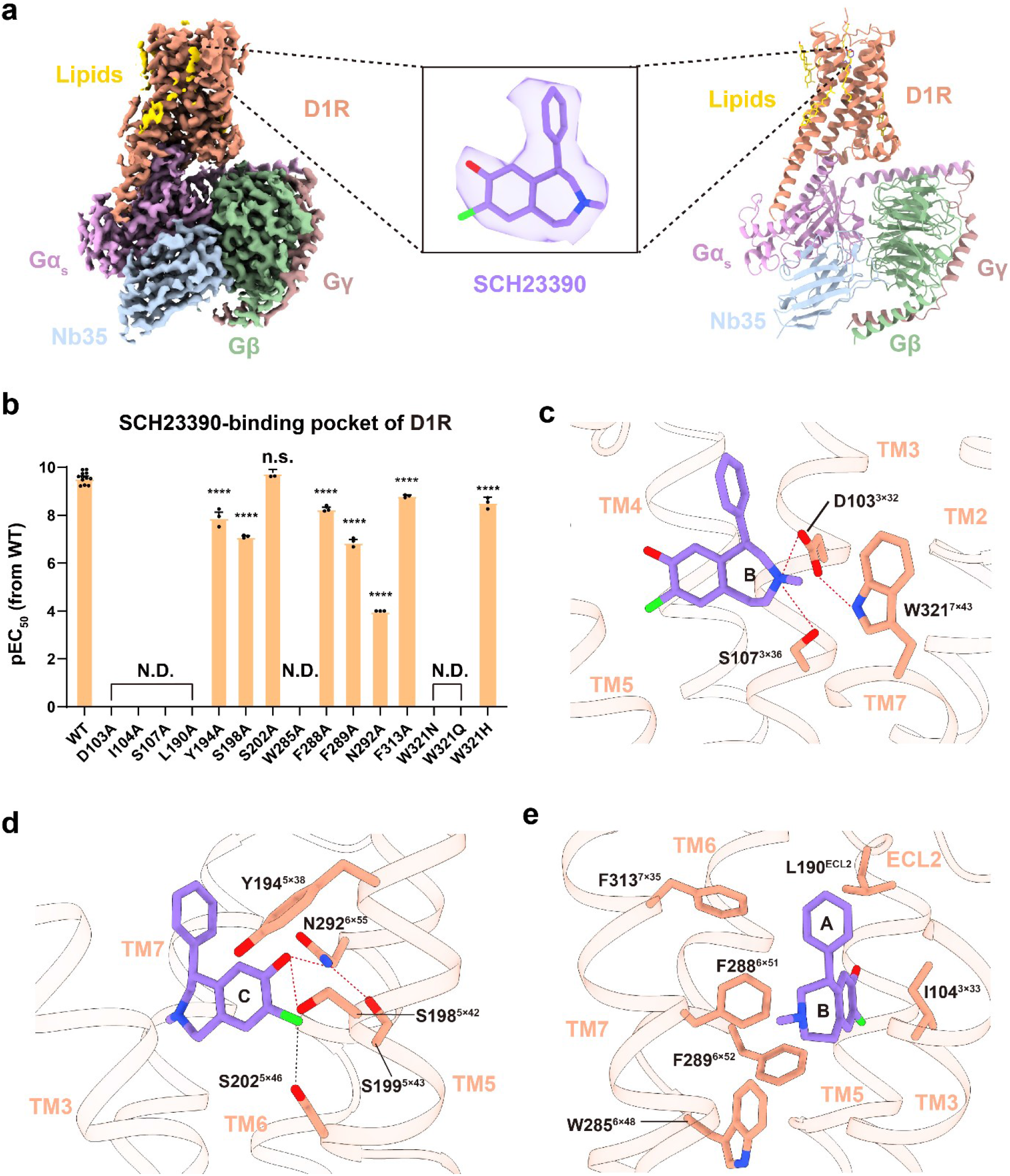
Structural basis of SCH23390 recognition by D1R. **a,** Cryo-EM density map and cartoon representation of the SCH23390-D1R-G_s_ complex. The inset shows SCH23390 and its cryo-EM density. D1R, Gα_s_, Gβ, Gγ, Nb35 and ordered lipids are colored separately. **b,** Potencies of SCH23390 at WT D1R and the indicated binding-pocket mutants, measured using the HTRF cAMP assay in HEK293 cells. Each point represents an independent experiment; data are mean ± S.E.M. from three to nine independent experiments (WT, *n* = 9; mutants, *n* = 3), each performed in triplicate. N.D., not determined, denotes either no response or a response for which reliable parameter estimates could not be established over the tested concentration range. For variants with quantifiable responses, *P* values were determined by one-way ANOVA with Dunnett’s multiple-comparisons test versus WT. \*\*\*\**P* < 0.0001; n.s., not significant. **c-d,** Two views of the polar interaction network anchoring SCH23390 in the orthosteric pocket. Red dashed lines indicate polar interactions involving D103^3.32^, S107^3.36^, W321^7.43^ (**c**) and Y194^5.38^, S198^5.42^, S199^5.43^, S202^5.46^, N292^6.55^ (**d**). **e,** Hydrophobic contacts between SCH23390 and residues from TM3, TM5, TM6, TM7 and ECL2.

Although SCH23390 adopts a binding pose highly similar to that of benzazepine agonists, its recognition within the catechol-binding region is distinct from that of ligands containing a catechol pharmacophore. Canonical catechol agonists, such as dopamine and the SKF-series compounds, engage an extensive hydrogen-bonding network through both the *meta*- and *para*-hydroxyl groups, serving as a key determinant of receptor activation^4,21^. The *meta*-hydroxyl of catechol group typically engages S198^5×42^ and S202^5×46^, whereas the *para*-hydroxyl primarily interacts with N292^6×55^ and, indirectly, S199^5×43^ (Extended Data Fig. 5b). However, in SCH23390, the *meta*-hydroxyl is replaced by a chloride substituent, leaving only the *para*-hydroxyl available for polar interactions. Consequently, SCH23390 forms close polar contacts through its *para*-hydroxyl with N292^6×55^ and S199^5×43^, along with a weaker interaction with S198^5×42^, while the chloride substituent instead contributes an additional van der Waals contact (Fig. 3d). Consistent with these observations, mutations disrupting the conserved anchor D103^3×32^ or the surrounding polar network, including S107^3×36^, N292^6×55^, and W321^7×43^, markedly reduced SCH23390-mediated signaling, whereas alanine substitution of S198^5×42^ or the adjacent residue Y194^5×38^ produced comparatively modest effects (Fig. 3b, Extended Data Fig. 4b and Supplementary Table 4). In addition, SCH23390 is further stabilized by hydrophobic interactions with surrounding residues including I104^3×33^, L190^ECL2^, W285^6×48^, F289^6×52^, and W321^7×43^ (Fig. 3e). Notably, SCH23390 carries a methyl group on its azepine ring analogous to that of the partial/biased agonist SKF83959, whose interaction with W321^7×43^ has been proposed as a critical determinant of partial or G protein-biased agonism of D1R^28^. Consistently, mutations at most of the residues contributing primarily hydrophobic contacts produced significantly reduced effects on D1R signaling (Fig. 3b, Extended Data Fig. 4b and Supplementary Table 4). Thus, these findings indicate that SCH23390 recognition is dominated by conserved polar interactions characteristic of D1R agonists, whereas the loss of one catechol hydroxyl reshapes the hydrogen-bonding network without altering the overall agonist-like binding mode.

### Conserved inverse agonism shared by D1R and D2R

Comparison of the LE300-bound inactive structure with the previously reported SKF81297-bound active D1R structure provides insight into how inverse agonist binding is translated into receptor inactivation. Although both ligands engage the conserved D103^3×32^-S107^3×36^-W321^7×43^ polar network, LE300 stabilizes a distinct interaction geometry that redirects its indole ring into the hydrophobic cavity adjacent to the conserved toggle switch W285^6×48^ (Figs. 4a-b). Unlike SKF81297, which leaves this cavity unoccupied, LE300 sterically restricts the rotation of W285^6×48^ into its active-state rotamer. This restraint prevents the coordinated rearrangement of the conserved PIF, sodium ion pocket, NPxxY, and DRY motifs, as well as the outward shift of TM3 toward TM2, thereby blocking the outward displacement of TM6 and inward movement of TM7 required to form the intracellular cavity for G protein coupling (Extended Data Figs. 3a and 6a). In addition to restraining the toggle switch, the LE300-bound structure reveals restoration of a polar interaction between T^1×46^ and S^7×47^ adjacent to the sodium ion pocket. This interaction is absent in active D1R structures and likely contributes to stabilization of the inactive sodium ion pocket conformation. Consistent with this structural observation, alanine substitution of either residue (T^1×46^A or S^7×47^A) significantly enhanced SCH23390-induced D1R activation (Extended Data Fig. 6b), supporting a role for this interaction in maintaining receptor inactivity.

**Fig. 4.**
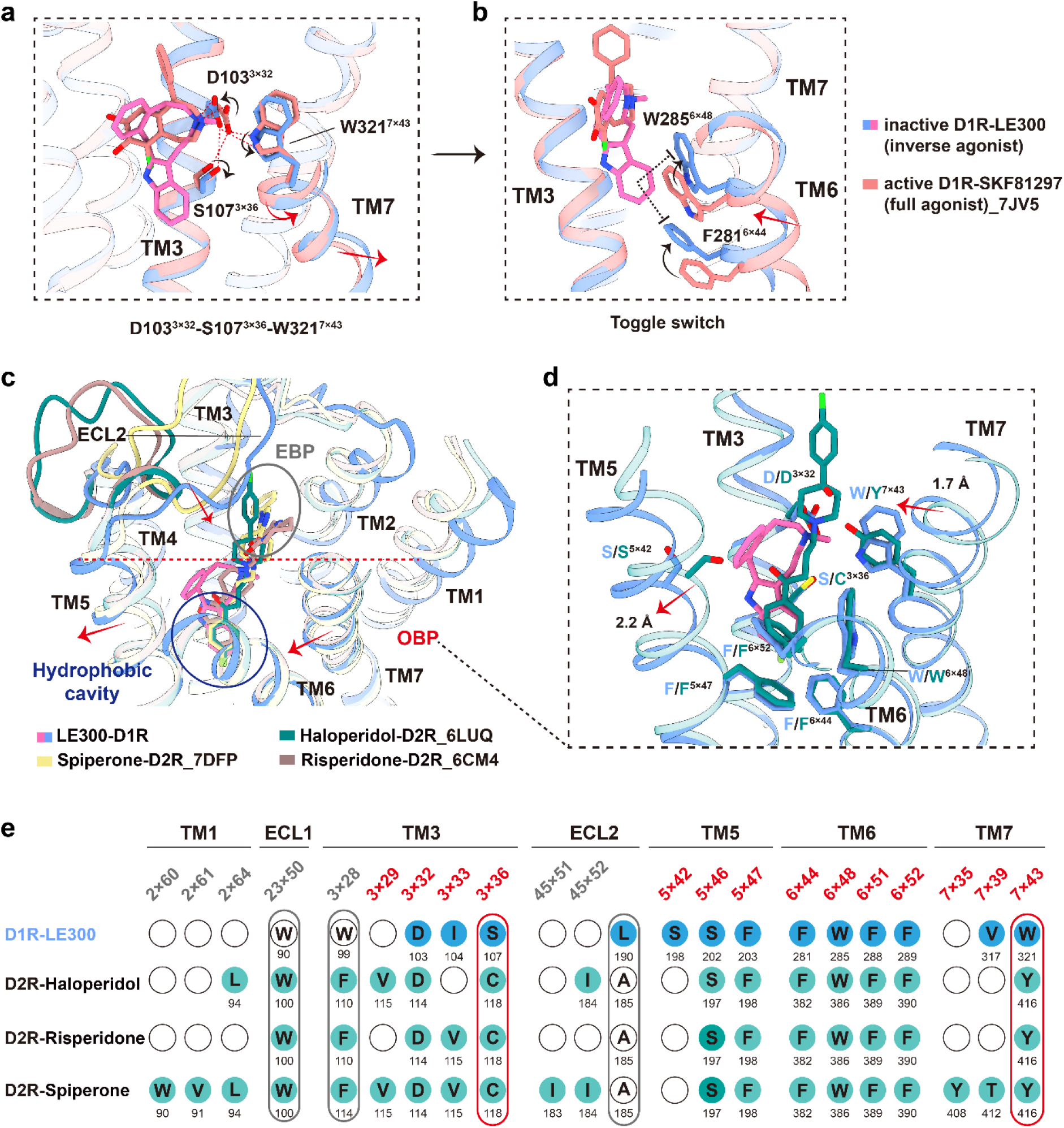
Conserved inverse agonism shared by D1R and D2R. **a-b,** Structural comparison of LE300-bound inactive D1R with the full-agonist SKF81297-D1R-G_s_ complex (PDB 7JV5), showing rearrangements of the D103^3.32^-S107^3.36^-W321^7.43^ polar network (**a**) and the W285^6.48^/F281^6.44^ toggle-switch region (**b**). Red dashed lines indicate polar interactions, and arrows indicate relative side-chain (black) and helical (red) displacements. **c,** Side-view superposition of LE300-bound D1R with inactive D2R structures bound to haloperidol (PDB 6LUQ), risperidone (PDB 6CM4) or spiperone (PDB 7DFP). The OBP and EBP are indicated. **d,** Close-up comparison of the D1R and D2R inverse-agonist binding pockets, highlighting ligand displacement and conserved and non-conserved positions in TM3, TM5 and TM7. The ellipse marks a predicted steric clash; dashed lines show measured separations and arrows indicate local conformational differences (black for side-chains; red for helices). **e,** Sequence alignment of ligand-contacting positions in LE300-bound D1R and the three inactive D2R-inverse agonist complexes. Filled circles denote ligand-contacting residues and open circles denote positions without a detected contact; outlined columns highlight subtype-selective positions. Red and gray labels indicate non-conserved positions in the OBP and EBP, respectively.

Functional characterization further showed that LE300 also exhibited inverse agonist activity at D2R under our assay conditions (Extended Data Fig. 6c and Supplementary Table 1). To determine whether this mechanism is unique to D1R or reflects a more general principle of DR inverse agonism, we compared the LE300-bound structure with previously reported inactive D2R structures bound to the inverse agonists haloperidol, risperidone, and spiperone^29–31^ (Figs. 4c-d). Despite substantial differences in ligand chemistry and receptor subtype, all of these inverse agonists occupy a hydrophobic cavity adjacent to W^6×48^ with a nearly identical residue composition that is not engaged by agonists, thereby stabilizing the inactive conformation of the toggle switch (Fig. 4c and Extended Data Fig. 6d). These observations suggest that direct stabilization of the W^6×48^-centered activation switch represents a conserved mechanism of inverse agonism across DRs.

Despite converging on the same inhibitory mechanism, D1R and D2R position their inverse agonists to engage the hydrophobic chamber through distinct recognition modes. Although several residues lining the OBP are conserved between the two receptor subtypes, subtype-specific substitutions reshape both the OBP and the extended binding pocket (EBP). In particular, replacement of Y^7×43^ and C^3×36^ in D2R with W^7×43^ and S^3×36^ in D1R reorganizes the conserved polar interaction network surrounding the ligand-binding site. In addition, the extracellular end of TM5 in D2R is displaced inward by approximately 2 Å relative to D1R, measured at the Cα atom of the conserved S^5×42^ residue, whereas the extracellular end of TM7 in D1R shifts inward by 1.7 Å relative to D2R at D/S^7×36^, resulting in a comparatively narrower extracellular vestibule (Fig. 4e). Together with additional non-conserved residues along TM2 and TM3 and distinct ECL2 conformations, these structural differences generate receptor subtype-specific ligand recognition environments (Fig. 4c-e and Extended Data Fig. 6e). Consequently, whereas D2R inverse agonists extend into the EBP to position their hydrophobic substituents toward the conserved W^6×48^-associated cavity, LE300 remains largely confined within the OBP while engaging the same inhibitory cavity (Fig. 4c).

### Progressive conformational changes underlying graded agonism at D1R

Having established the mechanism of inverse agonism of D1R, we next sought to determine how ligands spanning a range of efficacies generate graded receptor activation. SCH23390 and SKF83959 differ from the full agonist SKF81297 by only subtle chemical modifications, yet display markedly reduced efficacies (Figs. 1a, c). Structural comparison of the corresponding D1R-G_s_ complexes revealed that these ligands stabilize progressively different active conformations^4^. Most notably, the conserved toggle switch W285^6×48^ undergoes a graded downward swing that closely parallels ligand efficacy, with SCH23390, SKF83959, and SKF81297 occupying increasingly active conformations (Fig. 5a). These observations suggest that partial agonism arises from incomplete engagement of the conserved activation pathway.

**Fig. 5.**
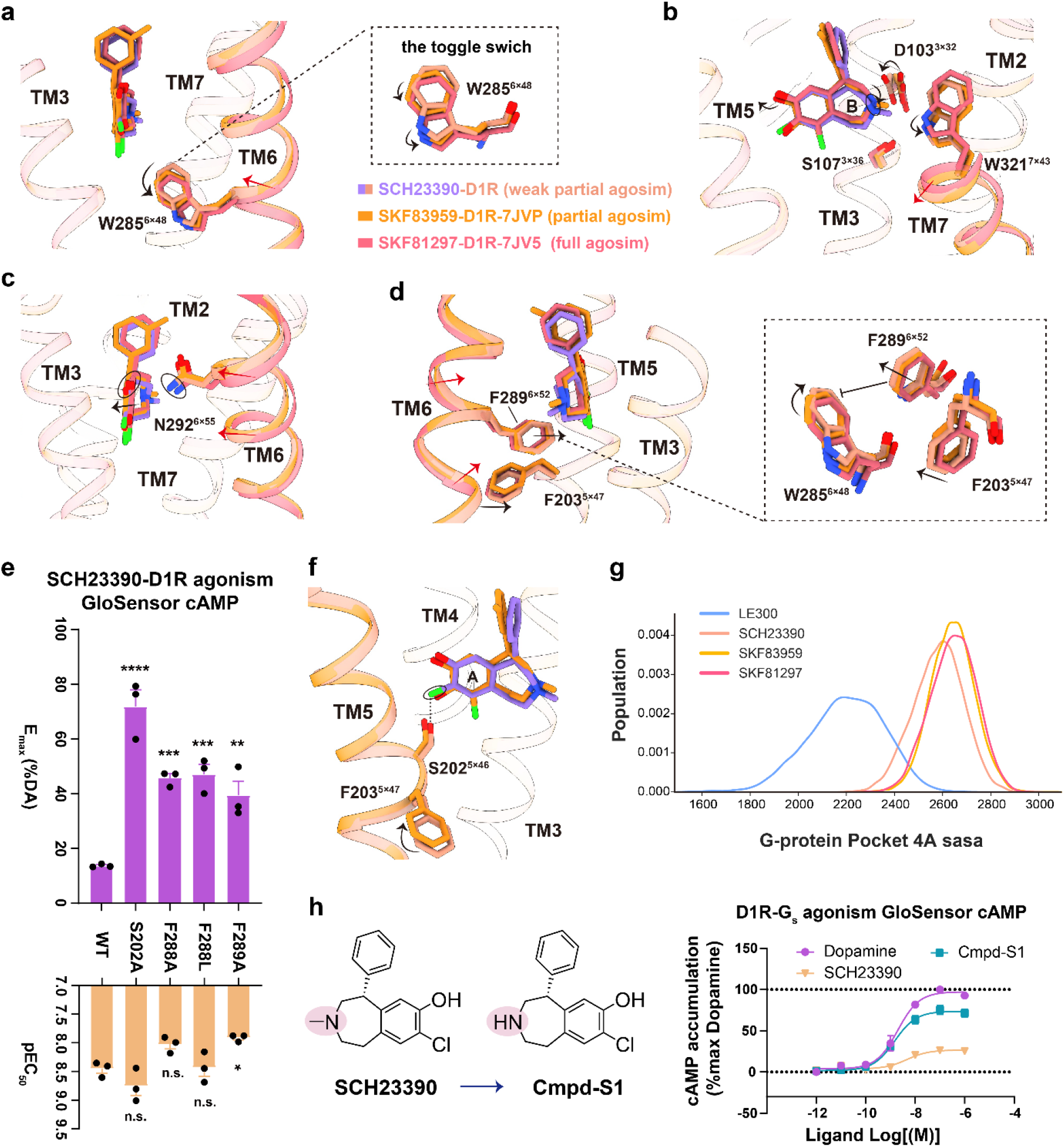
Structural determinants of graded agonism at D1R. **a,** Ligand-dependent rearrangements of the toggle switch W285^6.48^ in SCH23390, SKF83959 and SKF81297-bound D1R. Arrows indicate relative side-chain and TM6 displacements. **b,** Structural comparison of the D103^3.32^-S107^3.36^-W321^7.43^ polar network in SCH23390, SKF83959 and SKF81297-bound D1R. The superposition is followed by individual views of each ligand-receptor complex. Red dashed lines indicate polar interactions; black and red arrows indicate relative side-chain and helical displacements, respectively. The arrow between individual structures denotes increasing ligand efficacy. SKF83959 and SKF81297-bound structures correspond to PDB 7JVP and 7JV5, respectively. **c,** Ligand-dependent contacts with N292^6.55^ and associated movements of the extracellular portion of TM6 in SCH23390, SKF83959 and SKF81297-bound D1R. Arrows indicate relative ligand (black) and helical (red) displacements. **d,** Ligand-dependent packing of the F203^5.47^-F289^6.52^ aromatic motif and W285^6.48^ in SCH23390, SKF83959 and SKF81297-bound D1R. Red arrows indicate TM6 movements and black arrows indicate side-chain rearrangements. **e,** Potency and maximal efficacy of SCH23390 at WT D1R and the S202^5.46^, F288^6.51^ and F289^6.52^ mutants, measured using the GloSensor cAMP assay in HEK293 cells. Potency is expressed as pEC_50_ (-log_10_[EC_50_ (M)]), and Emax is expressed as a percentage of the maximal dopamine response at the corresponding receptor construct. Data are mean ± S.E.M. from three independent experiments (*n* = 3), each performed in triplicate. *P* values were determined by one-way ANOVA with Dunnett’s multiple-comparisons test versus WT. \**P* < 0.05, \*\**P* < 0.01, \*\*\**P* < 0.001 and \*\*\*\**P* < 0.0001; n.s., not significant. **f,** Structural comparison of SCH23390- and SKF83959-bound D1R around S202^5.46^. The dashed line indicates the chlorine-mediated SCH23390-S202^5.46^ interaction; arrows indicate relative ligand, side-chain and TM5 displacements. **g,** Distributions of the solvent-accessible surface area (SASA) of the intracellular G-protein-binding pocket in molecular dynamics simulations of LE300, SCH23390, SKF83959 and SKF81297-bound D1R, calculated using a 4-Å probe. **h,** Chemical structures of SCH23390 and its N-demethylated analogue Cmpd-S1 (Compound-S1), and concentration-response curves for dopamine, SCH23390 and Cmpd-S1 measured using the GloSensor cAMP assay in HEK293 cells expressing WT D1R. Responses were normalized to the maximal dopamine response (100%), with apo basal activity defined as 0%. Cmpd**-**S1 showed higher maximal efficacy than SCH23390. Data are mean ± S.E.M. from three independent experiments (*n* = 3), each performed in triplicate.

Comparison of the SCH23390-, SKF83959-, and SKF81297-bound structures further revealed the molecular determinants underlying this efficacy continuum. Relative to SKF81297, the additional methyl group present on the benzazepine ring of both SKF83959 and SCH23390 reduces engagement of the D103^3×32^-S107^3×36^-W321^7×43^ polar network and redirects the benzazepine scaffold slightly away from TM6 (Fig. 5b and Extended Data Fig. 7a). This repositioning facilitates a more pronounced inward movement of the extracellular portion of TM6, stabilized by a polar interaction between N292^6×55^ and the *para*-hydroxyl group of the ligand (Fig. 5c). This TM6 rearrangement in turn drives a conformational change in the adjacent F203^5×47^-F289^6×52^ aromatic motif that sterically precludes the downward swing of the toggle-switch residue W285^6×48^ required for full activation (Fig. 5d). Consistent with this model, alanine substitution of F289^6×52^ and its interaction residue F288^6×51^ enhanced the efficacy of both SCH23390 and SKF83959, while exerting comparatively smaller effects on dopamine-mediated activation (Fig. 5e and Extended Data Fig. 7b).

Unlike SKF83959, SCH23390 lacks the *meta*-hydroxyl group of the catechol scaffold, which is replaced by a chlorine substituent. Structural comparison of the two complexes revealed that this chlorine establishes a direct interaction with S202^5×46^, restricting the clockwise rotation of TM5 around F203^5×47^ (Fig. 5f). As a result, the F203^5×47^-F289^6×52^ motif undergoes a smaller transition toward its active-state conformation, further limiting the downward swing of the toggle switch W285^6×48^ beyond the effect imposed by the benzazepine methyl group. Consistent with this mechanism, alanine substitution of S202^5×46^ markedly increased the efficacy of SCH23390, demonstrating that this chlorine-mediated interaction constitutes a second, additive structural determinant of the reduced efficacy of SCH23390 (Fig. 5e and Extended Data Fig. 7b and Supplementary Table 5).

To corroborate this model, we performed molecular dynamics simulations of D1R bound to LE300, SCH23390, SKF83959, and SKF81297. Analysis of the intracellular G_s_ protein-binding cavity revealed a progressive expansion in both pocket size and conformational stability that closely tracked ligand efficacy, following the rank order SKF81297 > SKF83959 > SCH23390 > LE300 (Fig. 5g and Supplementary Table 6). In parallel, measurement of the distance between the *meta*-hydroxyl (or, in the case of SCH23390, the chlorine substituent) and S202^5×46^ across the three catechol-scaffold ligands revealed the rank order SKF81297 > SKF83959 > SCH23390, with the shortest distance observed for the SCH23390-S202^5×46^ interaction (Extended Data Fig. 7c and Supplementary Table 6). This trend is consistent with the structural model above, in which direct engagement of the chlorine substituent with S202^5×46^ restricts the active-state rotation of TM5 and thereby imposes an additional constraint on receptor activation specific to SCH23390.

Guided by the structural analysis, we designed Cmpd-S1 by removing the benzazepine methyl group from SCH23390 while preserving the remainder of the scaffold (Fig. 5h, Extended Data Fig. 8 and Supplementary Figs. 1-3). Remarkably, this single modification converted SCH23390 from a weak partial agonist into a high-efficacy partial agonist without altering its potency (Fig. 5h and Supplementary Table 1). The restored efficacy is consistent with re-establishment of the optimal D103^3×32^-S107^3×36^-W321^7×43^ interaction network and relief of the downstream conformational constraint imposed by the chlorine substituent on TM5, thereby enabling more complete propagation of the conformational changes leading to activation of W285^6×48^. These findings demonstrate that graded agonism at D1R is governed by progressive engagement of a conserved activation pathway.

## Discussion

Understanding how ligands encode distinct signaling efficacies remains a fundamental challenge in GPCR biology. While decades of biochemical, spectroscopic and computational studies have established that GPCRs populate dynamic conformational ensembles rather than simply inactive and active states, the structural basis by which these conformations generate graded signaling outputs has remained incompletely understood. Structural studies have largely captured the endpoints of receptor activation, whereas intermediate states have been inferred primarily from NMR, DEER spectroscopy and molecular dynamics simulations^32–35^. More recently, cryo-EM studies of several GPCRs have revealed subtle structural differences associated with ligands of different efficacies^4,36,37^. Nevertheless, these observations have generally represented isolated snapshots, and a structural framework connecting ligand chemistry, receptor conformation and signaling efficacy has remained lacking. In this study, we establish an experimentally connected activation landscape for D1R spanning inverse agonism, weak partial agonism, partial agonism and full agonism, demonstrating that graded GPCR activation is achieved through progressive engagement of a conserved activation pathway rather than discrete transitions between inactive and active receptor states.

A key finding emerging from this work is that ligand efficacy is determined by the extent to which ligands drive progression along a conserved activation trajectory. Our analyses demonstrate that ligands across the efficacy spectrum engage the same activation mechanism but differ in the efficiency with which conformational changes propagate from the OBP to the intracellular G protein-binding interface. Consequently, partial agonists do not activate fundamentally different receptor conformations but instead occupy intermediate positions along a common activation pathway. Central to this activation landscape is the highly conserved W^6×48^ toggle switch. Rather than functioning as a binary molecular switch, our structures and simulations indicate that W^6.48^ behaves as a continuously tunable conformational hub whose degree of activation closely parallels ligand efficacy, thereby linking intermediate receptor conformations with quantitatively distinct signaling efficacies. Progressive engagement of the D^3×32^-S^3×36^-W^7×43^ interaction network, together with coordinated rearrangements of TM5 and the adjacent F^5×47^-F^6×52^ aromatic motif, incrementally promotes W^6×48^ activation and expansion of the intracellular G protein-binding cavity.

Our findings also provide a molecular explanation for how chemically subtle ligand modifications generate disproportionately large pharmacological consequences. Rather than being determined by a single interaction, ligand efficacy emerges from the cumulative energetic contributions of multiple ligand-receptor contacts that collectively regulate propagation through the conserved activation pathway. This model is directly validated by the successful conversion of SCH23390 into a high-efficacy agonist through removal of a single methyl substituent. Beyond confirming the proposed mechanism, this prospective redesign demonstrates that efficacy can be predictably engineered through rational manipulation of allosteric signal propagation rather than empirical optimization.

Comparison with the D2R further reveals an unexpected combination of mechanistic conservation and structural divergence. Although inverse agonists occupy distinct binding environments in D1R and D2R, both receptors ultimately suppress activation by stabilizing the conserved W^6×48^-centered activation pathway. These observations suggest that closely related dopamine receptor subtypes have evolved distinct ligand-recognition mechanisms while converging on a common energetic bottleneck governing receptor activation. Such separation between ligand recognition and efficacy encoding may represent a general strategy by which homologous GPCRs achieve pharmacological diversity without altering their conserved activation machinery.

Beyond providing mechanistic insight into D1R pharmacology, our findings have broader implications for GPCR biology. Conserved activation microswitches, including the toggle switch, PIF motif, sodium pocket and NPxxY motif, are widely shared across class A GPCRs. Previous structural studies have primarily described these motifs as binary activation switches. Our data instead suggest that these microswitches can be engaged progressively, allowing receptors to populate multiple intermediate conformations that encode distinct signaling efficacies. This model offers a structural explanation for the widespread phenomenon of graded agonism observed across GPCR families and may represent a general principle governing efficacy modulation in class A GPCRs.

Finally, these findings have important implications for GPCR drug discovery. Therapeutic optimization has traditionally focused on improving ligand affinity, potency and subtype selectivity, whereas ligand efficacy has remained difficult to control rationally. Our results demonstrate that efficacy can be predictably tuned by selectively modulating propagation through conserved activation networks, offering a route to optimize therapeutic benefit while preserving physiological signaling balance. More broadly, this work linking ligand chemistry to graded receptor activation and provides a molecular foundation for the rational design of efficacy-tuned therapeutics with precise control over signaling output at DRs and other class A GPCRs.

## Acknowledgements

This work is supported by the following fundings: The National Natural Science Foundation of China (32130022, 82495184 to H.E.X.; 82121005 to H.E.X. and X.X.; 32401002 to Y.Z.; 82330113 to X.X.; 82304579 to S.G.); the National Key R&D Program of China (2022YFC2703105, 2022YFE0203600 to H.E.X.; 2024YFA1307504 to Y.Z.); Shanghai Municipal Science and Technology Major Project (2019SHZDZX02 to H.E.X.); Strategic Priority Research Program of the Chinese Academy of Sciences (XDB0830000, XDA0530000, XDB37030103 to H.E.X); the Natural Science Foundation of Shanghai, China (23ZR1475300 to Y.Z.); the Sailing Program of Shanghai Venus Project (23YF1456700 to Y.Z.); the HE Research Fellowship (HERF2025011 to Y.Z.). The cryo-EM data of this study were collected at the Advanced Center for Electron Microscopy, Shanghai Institute of Materia Medica (SIMM). We thank all the staffs at the center for their assistance in cryo-EM data collection. We also thank Shanghai Frontiers Science Center of Cellular Homeostasis and Human Diseases of Shanghai Jiao Tong University, School of Medicine for the technical support.

## Author contributions

Y.W.Z. and H.E.X. initiated the project. X.Y.Z. optimized the constructs of D1R, prepared the protein samples of LE300-D1R and SCH23390-bound D1R-G_s_-Nb35 complexes for cryo-EM studies, screened the cryo-EM grids, performed cryo-EM data acquisition, prepared the Figures and Tables, and participated in manuscript editing; Y.Z., J.S. and S.G. performed the GloSensor and HTRF cAMP functional assays and participated in figure preparation. H.L. conducted the structure determination and model coordinates building, and participated in manuscript editing and Figures preparation; L.F. participated in protein sample preparation and optimization; J.H. and X.H. performed the MD simulations and participated in figure preparation. T.L. and J.Z. synthesized Cmpd-S1 and wrote the chemical part of this manuscript; R.L. and M.L. participated in functional data analysis; W.H. assisted in cryo-EM data collection; X.C. supervised J.H. and X.H. in MD simulation studies; S.W. supervised L.F. in protein sample preparation and optimization; S.G. and X.X. supervised Y.Z. and J.S. in functional assays; Y.W.Z. wrote the manuscript with the inputs from the others; Y.W.Z. and H.E.X. conceived and supervised the project.

## Data availability

The cryo-EM density maps have been deposited in the Electron Microscopy Data Bank under accession numbers EMD-82510 (D1R bound to LE300) and EMD-82511 (D1R-G_s_-Nb35 complex bound to SCH23390). The corresponding atomic coordinates have been deposited in the Worldwide Protein Data Bank (wwPDB) under accession numbers 44CW and 44CX, respectively. All other data supporting the findings of this study are available in the main text and supplementary Figures and Tables.

## Materials and Methods

### Cell lines

*Spodoptera frugiperda* (Sf9 cells, Expression Systems) were maintained in ESF 921 serum-free medium at 27 °C with agitation at 120 rpm. Sf9 cells were used for baculovirus production and expression of the receptor complexes. Escherichia coli strains used for Nb35 and anti-BRIL Fab production are specified below. Human embryonic kidney 293 (HEK293) cells were used in functional activity assays.

### Constructs

The coding sequence of full-length human D1R was cloned into pFastBac. For preparation of the SCH23390–D1R–miniG_s_ complex, the receptor carried an N-terminal prolactin signal peptide followed by a FLAG epitope and a β_2_-adrenergic receptor N-terminal fragment (BN). A TEV-protease site separated BN from D1R, and a His8 tag was placed at the receptor C terminus. The dominant-negative miniG_αs_ construct lacked the α-helical domain and contained G226A and A366S substitutions^38–40^. Human miniG_αs_, rat G_β_1__ and bovine G_γ_2__ were cloned separately in pFastBac, with an N-terminal His₈ tag on Gβ_1_. This design followed the established D1R-miniGs strategy^4^.

For the LE300-bound receptor, BRIL (thermostabilized apocytochrome b₅₆_2_RIL) replaced the central portion of D1R intracellular loop 3 (ICL3). D1R residues 1-223 and 267-446 were connected to BRIL through short linkers^41^. The construct contained S110K, I111A, L112W, L274A and M278A substitutions, the same N-terminal signal peptide-FLAG-BN module and TEV site described above, and a C-terminal His₈ tag. Nanobody-35 (Nb35) with a C-terminal His8 tag, was expressed in the periplasm of *E. coli* strain BL21^42^.

The anti-BRIL Fab heavy and light-chain sequences encoded separately in pFastBac, with the GP67 signaling peptide was inserted at the N-terminus, and a TEV-His8 tag was fused to the C-terminus.

### Preparation of Nb35

Nb35 was expressed in the periplasm of *E. coli* BL21. Cells were grown in Terrific Broth supplemented with 0.1% (w/v) glucose, 2 mM MgCl_2_ and 100 µg mL⁻¹ ampicillin at 37 °C to an OD₆₀₀ of approximately 1.0. Expression was induced with 1 mM IPTG for 4.5 h at 37 °C. Harvested cells were resuspended in ice-cold 50 mM Tris-HCl, pH 8.0, 12.5 mM EDTA and 0.125 M sucrose. After removal of cell debris, Nb35 was purified by immobilized-metal affinity chromatography and size-exclusion chromatography on a HiLoad 16/600 Superdex 75 column. Monodisperse fractions were pooled, concentrated to approximately 2 mg mL⁻¹, flash-frozen in liquid nitrogen and stored at −80 °C.

### Expression and purification of anti-BRIL Fab

Nb35 was expressed in the Sf9 cells. Cell supernatant was collected 48 h after infection at 27 °C. The anti-BRIL Fab stocks were prepared by methods previously described (as scfv16^43^), including nickel affinity chromatography followed by size exclusion chromatography on a HiLoad 16/600 Superdex 75 column (Cytiva). Monodisperse Fab was concentrated to approximately 3 mg mL⁻¹, flash-frozen and stored at −80 °C.

### Expression and purification of the SCH23390-D1R-miniGs-Nb35 complex

Recombinant baculoviruses encoding D1R, miniG_αs_, G_β_1__ and G_γ_2__ were used to infect Sf9 cells at 3× 10⁶ cells mL⁻¹ at a virus ratio of 1:1:1:1. Cells were collected 48 h after infection at 27 °C and stored at −80 °C. Pellets from a 0.5-L culture were resuspended in 20 mM HEPES, pH 7.3, 50 mM NaCl, 5 mM CaCl_2_, 5 mM MgCl_2_, 0.3 mM TCEP and protease inhibitors. Membrane-associated complex formation was promoted by adding 5 µM SCH23390 and 25 mU mL⁻¹ apyrase, followed by incubation for 1.5 h at room temperature. Membranes were collected at 100,000 × g for 45 min.

The membrane pellet was solubilized for 2 h at 4 °C in 20 mM HEPES, pH 7.3, 100 mM NaCl, 25 mM imidazole, 5 mM CaCl_2_, 10% (v/v) glycerol, 0.3 mM TCEP, 0.5% (w/v) LMNG, 0.1% (w/v) CHS, 0.025% (w/v) digitonin, 1 µM SCH23390, 25 mU mL⁻¹ apyrase and 10 µg mL⁻¹ Nb35. Insoluble material was removed at 100,000 × g for 45 min, and the supernatant was incubated with pre-equilibrated anti-FLAG resin overnight at 4 °C. The resin was washed sequentially with 10 column volumes of 20 mM HEPES, pH 7.3, 100 mM NaCl, 0.3 mM TCEP and 1 µM SCH23390 containing first 0.05% LMNG and 0.01% CHS and then 0.01% LMNG and 0.002% CHS. Bound complex was eluted in the latter buffer supplemented with 0.025% digitonin and 200 µg mL⁻¹ FLAG peptide.

The eluate was concentrated in a 100-kDa molecular-weight-cutoff centrifugal device and separated on a Superdex 200 Increase 10/300 GL column equilibrated with 20 mM HEPES, pH 7.3, 100 mM NaCl, 0.00075% LMNG, 0.00025% GDN, 0.0002% CHS, 0.005% digitonin, 0.1 mM TCEP and 5 µM SCH23390. Fractions containing monodisperse SCH23390-D1R-miniGs-Nb35 complex were pooled and concentrated to approximately 8 mg mL⁻¹ for cryo-EM.

### Expression and purification of the LE300-D1R-anti-BRIL Fab complex

Sf9 cells at 3 × 10⁶ cells mL⁻¹ were infected with D1R–BRIL baculovirus and harvested after 48 h at 27 °C. A pellet corresponding to 0.5 L of culture was resuspended in 20 mM HEPES, pH 7.3, 100 mM NaCl, 5 mM CaCl_2_, 5 mM MgCl_2_ and protease inhibitors. Membranes were solubilized for 3 h at 4 °C with 0.5% (w/v) LMNG, 0.1% (w/v) CHS and 100 µM LE300. After centrifugation at 100,000 × g for 45 min, the supernatant was incubated with anti-FLAG resin for 3 h at 4 °C. Detergent was reduced stepwise by washing with 10 column volumes of 20 mM HEPES, pH 7.3, 100 mM NaCl and 10 µM LE300 containing first 0.05% LMNG and 0.01% CHS and then 0.01% LMNG and 0.002% CHS. Receptor was eluted with 20 mM HEPES, pH 7.3, 100 mM NaCl, 0.3 mM TCEP, 10 µM LE300, 0.025% digitonin, 0.01% LMNG, 0.002% CHS and 200 µg mL⁻¹ FLAG peptide.

Purified LE300-D1R-BRIL was mixed with a 1.2-fold molar excess of anti-BRIL Fab and incubated on ice for 1 h, following the receptor–Fab assembly strategy used for D2R–Fab3089 while retaining an anti-BRIL-specific Fab (Im et al., 2020). The sample was concentrated and subjected to size-exclusion chromatography on a Superdex 200 Increase 10/300 GL column in 20 mM HEPES, pH 7.3, 100 mM NaCl, 10 µM LE300, 0.00075% LMNG, 0.00025% GDN, 0.0002% CHS and 0.05% digitonin. Peak fractions containing the assembled complex were pooled and concentrated to approximately 9.5 mg mL⁻¹ for cryo-EM.

### Cryo-EM grid preparation and data collection

For each complex, 3 µL of purified sample at 8-9.5 mg mL⁻¹ was applied to a freshly glow-discharged holey Nitinol grid (M01 Au300 R1.2/1.3; Nanodim). Grids were vitrified in liquid ethane using a Vitrobot Mark IV and stored in liquid nitrogen. Images were collected on a 300-kV Titan Krios microscope at the Advanced Center for Electron Microscopy, Shanghai Institute of Materia Medica, Chinese Academy of Sciences. For the LE300-D1R-BRIL-anti-BRIL Fab complex, two datasets comprising 3,720 and 5,946 movies (9,666 in total) were recorded at a nominal magnification of 105,000×, a calibrated pixel size of 0.724 Å, a defocus range of −1.0 to −2.0 µm and a total exposure of 50 e⁻ Å⁻². For the SCH23390-D1R-miniGs-Nb35 complex, 4,220 movies were acquired at a nominal magnification of 81,000×, a calibrated pixel size of 0.81 Å, the same defocus range and a total exposure of 80 e⁻ Å⁻². Each movie was divided into 36 frames during motion correction^44^.

### Image processing and map reconstruction

For the LE300 dataset, dose-fractionated movies were corrected for motion correction by MotionCor2.1^45^. Contrast transfer function (CTF) parameters for micrograph were estimated by patch CTF estimation. Particle processing was performed in cryoSPARC v3.2^46^. Blob-based and template-assisted picking yielded 3,835,046 particles. Reference-free two-dimensional classification, ab initio reconstruction and two rounds of heterogeneous refinement reduced the dataset to 404,416 particles. Non-uniform and local refinement, followed by receptor-focused refinement with a soft mask, produced a 3.6 Å receptor map.

The SCH23390-D1R-miniG_s_-Nb35 dataset was processed using the RELION3^47^ workflow described for D1R signaling complexes. Movie stacks were motion-corrected and dose-weighted with MotionCor2.1^45^, and contrast-transfer-function parameters were determined with Gctf v1.06^48^. Auto-picking from 4,220 movies yielded 2,734,371 particles. Reference-free two-dimensional classification and maximum-likelihood three-dimensional classification were performed in RELION 3.0 with C1 symmetry. The selected 223,026 particles were subjected to three-dimensional auto-refinement, Bayesian polishing and final three-dimensional refinement, producing a 3.2-Å map at the gold-standard FSC = 0.143 criterion. Local resolution was calculated from the two half-maps using Bsoft.

### Model building and refinement

For the SCH23390-D1R-miniG_s_-b35 complex, the structure of reported D1R (PDB: 7JV5) was used as the initial template. The model was adjusted to the sequences and engineered miniG_s_ construct used here, and unsupported regions were omitted. For the LE300 complex, inactive D2R (PDB 7DFP) was used as the initial template. LE300 and SCH23390 were placed only after their densities were clearly resolved. Initial models were docked into the cryo-EM maps in UCSF Chimera and iteratively rebuilt in Coot^49^. Real-space and reciprocal-space refinement were performed in Phenix with stereochemical and secondary-structure restraints, and geometry was assessed with MolProbity. Overfitting was evaluated by refining each model against one half-map and comparing map-versus-model FSC curves against both half-maps and the combined map. Refinement statistics are provided in the corresponding supplementary table. Structural illustrations were prepared with ChimeraX and PyMOL (https://pymol.org/2/).

### HTRF cAMP Assay

The DRD1-WT gene was sub-cloned in vector pcDNA3.0, This construct have a N-terminal haemagglutinin signal peptide. Mutations were introduced by QuickChange PCR. Sequences of mutants were verified by DNA sequencing. HEK293 cells were cultured in 1 x DMEM supplemented with 10% (v/v) fetal bovine serum and incubate in 5% CO_2_ at 37 ℃. For transient transfection, approximately 2×10^6^ cells were mixed with 1 μg of plasmids in 200 μl of transfection buffer, and electroporation was performed with a Scientz-2C electroporation apparatus (Scientz Biotech, Ningbo, China). Transfected cells were seeded into 10-cm cell culture dishes and incubated overnight for 24 h. The cells were then trypsinized, resuspended in serum-free DMEM containing 0.1% BSA, and supplemented with IBMX at a final concentration of 0.5 mmol/L. Cells were plated into 384-well plates at a density of 2,000 cells per well, followed by the addition of 5 μL of agonists at varying concentrations. The plates were incubated at 37 °C in the dark for 30 min. Intracellular cAMP levels were tested by a LANCE Ultra cAMP kit (Revvity, TRF0264) and EnVision multiplate reader according to the manufacturer’s instructions.

### Glosensor™ cAMP Assay

HEK293 cells (2.5 × 10⁶) were co-transfected with 2 μg WT or mutant DRD1 plasmid and 0.5 μg GloSensor plasmid using a Scientz-2C electroporator. The transfected cells were seeded into white 96-well plates and cultured overnight for 24 h. The culture medium was then removed, and 50 μL per well of Glosensor substrate (composed of 88% CO_2_-independent medium, 10% fetal bovine serum, and 2% Glosensor™ cAMP Reagent stock solution (Progema, E1291)) was added. The plates were incubated at 37 °C in a cell culture incubator for 2 h, followed by incubation at room temperature in the dark for 1 h. Compounds at serially diluted concentrations were added at 25 μL per well. Finally, luminescence was measured using an Envision2104 multimode microplate reader.

### MD simulations

The simulation systems were derived from the D1R-SKF81297 (PDB ID: 7JV5), D1R-SKF83959 (PDB ID: 7JVQ), D1R-SCH23390 and D1R-LE300 complexes. G proteins were excluded from the system. The complexes were incorporated into a POPC lipid bilayer using the packmol-memgen software^50^, sized to accommodate the extended length of T of TM6 (see Appendix Table Sxx for details), and were surrounded by a 12 Å aqueous layer. The ionic strength was maintained at 0.15 mol/L NaCl, with counterions added to balance the system. The FF19SB, Lipid17, and GAFF2 force fields were employed for amino acids, lipids, and ligands, respectively^51–53^. Each system underwent energy minimization followed by heating and equilibration according to established protocols^54,55^. Three independent 500 ns production runs were conducted using pmemd.cuda in Amber22 under the NVT ensemble at 300 K and 1 atm^56^. Long-range electrostatic interactions were computed using the Particle Mesh Ewald method, while short-range electrostatic and van der Waals interactions used a 10 Å cutoff. The SHAKE algorithm and hydrogen mass repartitioning were applied to constrain hydrogen-containing bonds, allowing a timestep of 4 fs. CPPTRAJ was used to calculate distance. Molecular mechanics generalized Born surface area (MM-GBSA).

### Synthetic detail of Cmpd-S1

#### 3-Chloro-4-methoxyphenethylamine (1)

3-Chloro-4-methoxyphenylacetonitrile (5 g, 27.53 mmol) was dissolved in dry THF, and the solution was added dropwise with vigorous stirring under N_2_ atmosphere to a solution of 2.0 M BH_3_-DMS (68.83 mL, 137.65 mmol). The mixture was refluxed under N_2_ for 2 h, and 25 mL of MeOH was slowly added. The mixture was again refluxed for 0.5 h and then concentrated in vacuo. The residue was diluted with water, extracted by EA, washed with brine and dried over Na_2_SO_4_. The crude product was purified by silica gel column to give compound **1**, which was a yellow oil. HRMS(ESI): m/z 169.1099 (M−NH_2_)^+^.

#### 2-((3-chloro-4-methoxyphenethyl)amino)-1-phenylethan-1-ol (2)

Compound **1** (2 g, 10.77 mmol), acetic acid (647 mg, 10.77 mmol) and styrene oxide (1.94 g, 16.16 mmol) were dissolved in anhydrous CH_3_CN. The mixture was stirred under reflux for 16 h in an atmosphere of N_2_. It was then cooled and concentrated in vacuo. The residue was diluted with water, extracted by EA, washed with brine and dried over Na_2_SO_4_, filtered, and concentrated in vacuo. The residual oil was triturated with Et_2_O. The precipitate was filtered, washed with anhydrous Et_2_O until colorless, and dried in vacuo to yield compound **2** which was a white solid. HRMS(ESI): m/z 306.2389 (M+H)^+^.

#### tert-butyl (3-chloro-4-methoxyphenethyl)(2-hydroxy-2-phenylethyl)carbamate (3)

To a stirred solution of compound **2** (1 g, 3.27 mmol) in DCM was added di-tert-butyl dicarbonate (785 mg, 3.6 mmol) and Et_3_N (662 mg, 6.54 mmol). After overnight stirring, the reaction mixture was diluted with water, and extracted by DCM, washed with brine and dried over Na_2_SO_4_. The crude product was purified by silica gel column to give compound **3**, which was a yellow oil. HRMS(ESI): m/z 406.1755 (M+H)^+^.

#### 7- chloro-8-methoxy-1-phenyl-2,3,4,5-tetrahydro-1H-benzo[d]azepine (4)

To a solution of compound **3** (500 mg, 1.23 mmol) in trifluoroacetic acid was added dropwise sulfuric acid (18 M, 103 μL, 1.85 mmol). The reaction was refluxed for 3 h. After evaporating the trifluoroacetic acid under reduced pressure, the resulting crude mixture was poured into ice cold water. The solution was made alkaline with 40% NaOH and extracted with EA. The organic layers were washed with brine, dried over Na_2_SO_4_, filtered and concentrated under reduced pressure to give compound **4**, which was a pale solid. HRMS(ESI): m/z 288.1480 (M+H)^+^.

#### 8- chloro-5-phenyl-2,3,4,5-tetrahydro-1H-benzo[d]azepin-7-ol (5)

To a solution of compound **4** (300 mg, 1.04 mmol) in dry DCM was added BBr_3_ (2 M in DCM, 5.6 mL) in a −20 °C bath, the resulting mixture was stirred for 6 h at −20 °C. Once TLC analysis indicated that the reaction was complete, the resulting mixture was poured into cold water and extacted with DCM. The organic phase was collected and dried over Na_2_SO_4_, filtered, and concentrated under vacuum, which was purified by column chromatography to afford compound **5** as a white solid. ^1^H NMR (400 MHz, DMSO-*d*_6_) *δ* 7.36 (t, *J* = 7.5 Hz, 2H), 7.25 (t, *J* = 7.4 Hz, 1H), 7.17 −7.08 (m, 3H), 6.32 (s, 1H), 4.19 (d, *J* = 7.6 Hz, 1H), 3.26 (m, 1H), 3.12 (m, 1H), 2.88 (m, 2H), 2.75-2.62 (m, 2H).

Compounds **S1** and **S2** were obtained via chiral separation. Compounds **S1** and **S2** were isolated through CHIRALPAK AD-H (ADH0CE-CW056) column (0.46 cm I.D. × 25 cm L), mobile phase: methanol / diethylamine = 100/0.1 (v/v), flow rate: 1.0 mL/min, wavelength: UV 214 nm, temperature: 35°C, HPLC equipment: Shimadzu LC-20AT, Compound **S1** was eluted as the first peak before the other enantiomer Compound **S2** (Compound **S1** Ret. time: 3.868 min; Compound **S2** Ret. time: 5.060 min), Compounds **S1** and **S2** both have enantiomeric excess (ee) values of >98%.

---

**Extended Data Fig. 1.**
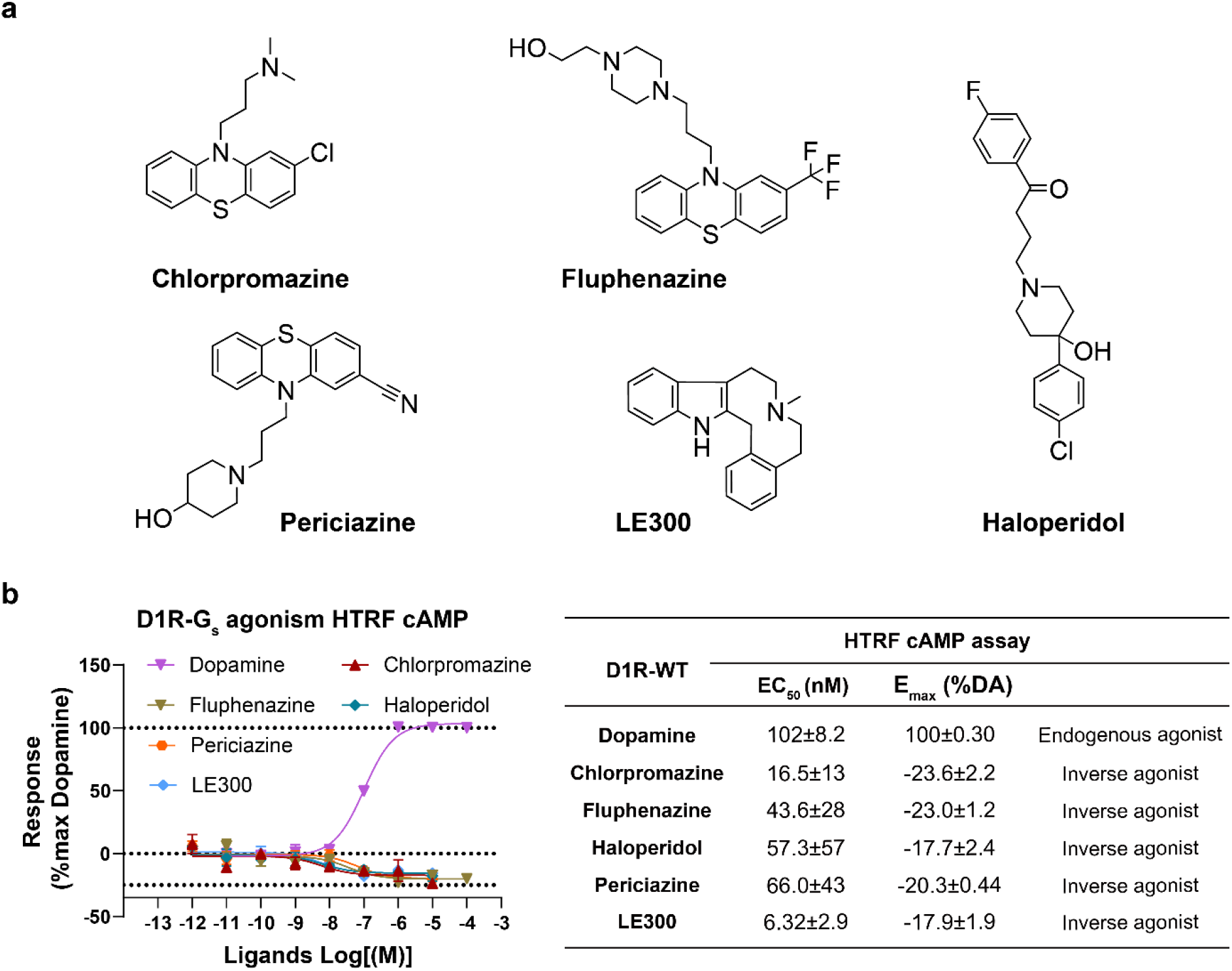
Functional screening of D1R inverse agonists. **a,** Chemical structures of chlorpromazine, fluphenazine, periciazine, haloperidol and LE300. **b,** Concentration-response curves for Chlorpromazine, Fluphenazine, Periciazine, Haloperidol and LE300 in HEK293 cells expressing the wild-type (WT) D1R, measured using the HTRF cAMP assay. Responses were normalized to the maximal dopamine response (100%), with apo basal activity defined as 0%. Data are mean ± S.E.M. from three independent experiments (*n* = 3), each performed in triplicate. Summary of EC_50_, Emax and pharmacological activity for these compounds shown in the right table. The five compounds are D1R inverse agonists. DA, dopamine.

**Extended Data Fig. 2.**
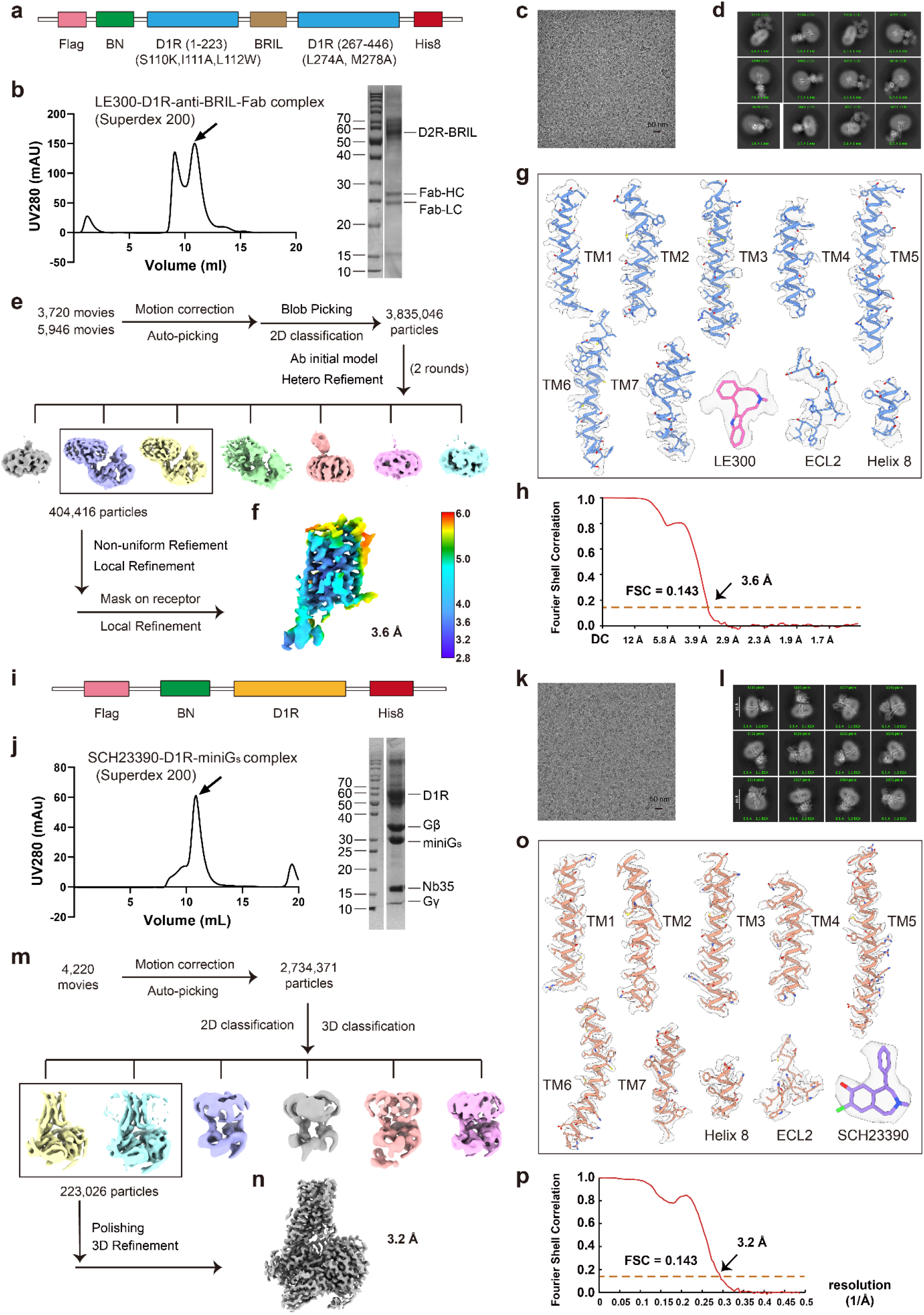
Sample preparation and cryo-EM of LE300-D1R-anti-BRIL-Fab complexes and SCH23390-D1R-G_s_ complexes. **a**, Cartoon models of the D1R constructs used in this study. **b,** Representative size exclusion chromatography (SEC) profiles and SDS-PAGE analysis of D1R-anti-BRIL-Fab bound to LE300. Experiment was repeated at least three times with similar results. **c-d,** Representative cryo-EM image from 9,666 movies (**c**) and 2D classification averages (**d**) of LE300-D1R-anti-BRIL-Fab. **e,** Cryo-EM data processing flowcharts of LE300-D1R-anti-BRIL-Fab by cryoSPARC 3.2. **f,** The global density map of LE300-D1R (mask on receptor) colored by local resolutions. **g,** The density maps of helices TM1-TM7 of transmembrane domain, extracellular loop ECL2 of D1R, and LE300 in LE300-D1R-anti-BRIL-Fab complex. **h,** The Fourier shell correlation (FSC) curves of LE300-D1R-anti-BRIL-Fab. The global resolution of the final processed density map estimated at the FSC = 0.143 is 3.6 Å. **i,** Cartoon models of the D1R constructs used in this study. **j,** Representative size exclusion chromatography (SEC) profiles and SDS-PAGE analysis of D1R-G_s_ complex bound to SCH23390. Experiment was repeated at least three times with similar results. **k-l,** Representative cryo-EM image from 4,220 movies (**k**) and 2D classification averages (**l**) of SCH23390-D1R-G_s_. **m,** Cryo-EM data processing flowcharts of SCH23390-D1R-G_s_ by Relion 3.0. **n,** The global density map of SCH23390-D1R-G_s_. **o,** The density maps of helices TM1-TM7 of transmembrane domain, extracellular loop ECL2 of D1R, and SCH23390 in SCH23390-D1R-G_s_ complex. **p,** The Fourier shell correlation (FSC) curves of l-SPD-D1R-G_s_. The global resolution of the final processed density map estimated at the FSC = 0.143 is 3.2 Å.

**Extended Data Fig. 3.**
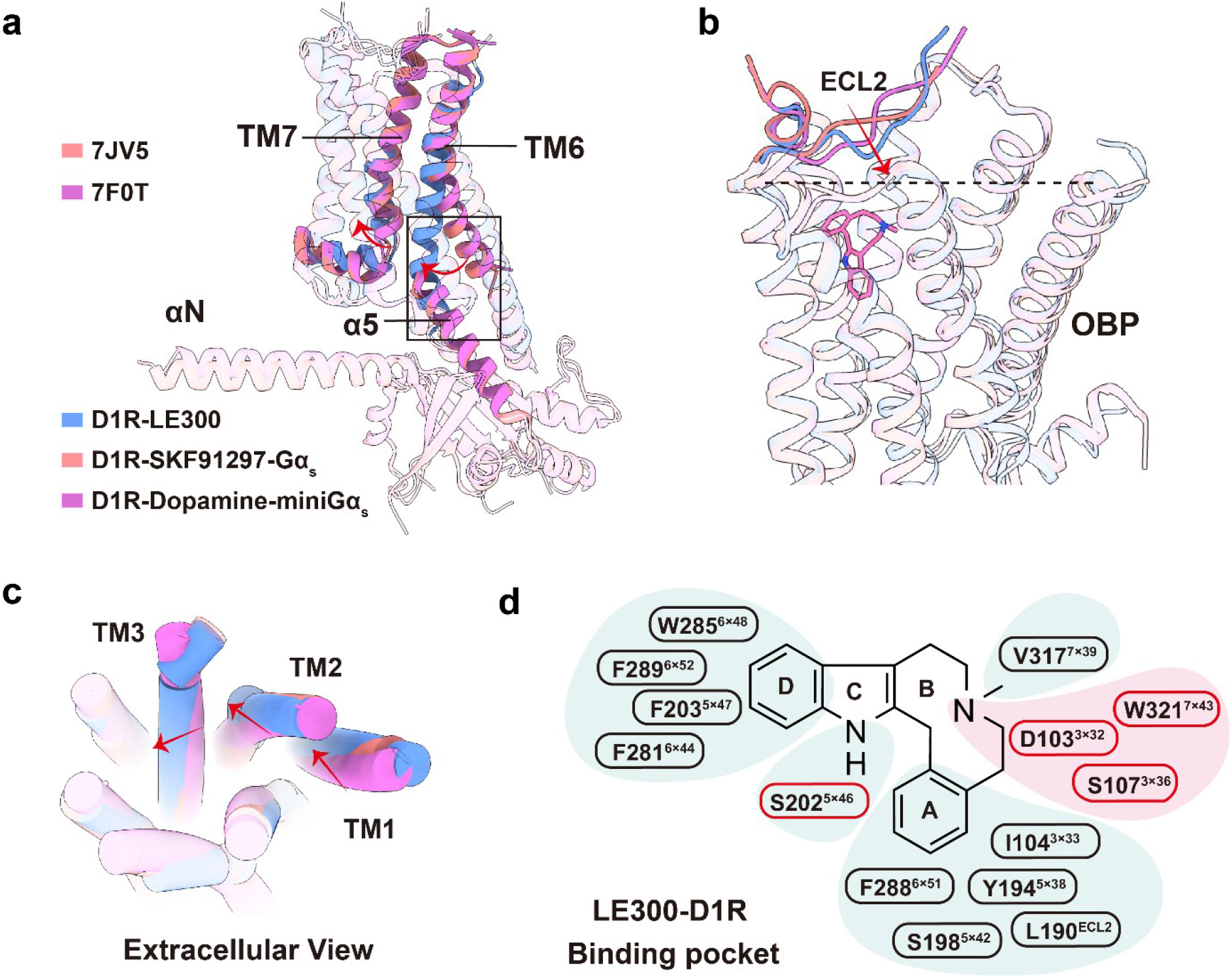
Additional conformational and ligand-binding features of the LE300-bound inactive D1R. **a,** Side-view superposition of LE300-bound inactive D1R with the SKF81297-D1R-G_αs_ complex (PDB 7JV5) and dopamine-D1R-miniG_αs_ complex (PDB 7F0T). **b,** Side-view comparison highlighting the inward displacement of ECL2 in the LE300-bound inactive state relative to the SKF8129 and dopamine-bound active states. **c,** Extracellular view of the same superposition, highlighting coordinated movements of ECL2 and the extracellular portions of TM1, TM2 and TM3 in the LE300-bound inactive state. Red arrows indicate relative displacements. **d,** Two-dimensional summary of residues contacting LE300. Polar and hydrophobic contact regions are indicated by pink and blue shading, respectively.

**Extended Data Fig. 4.**
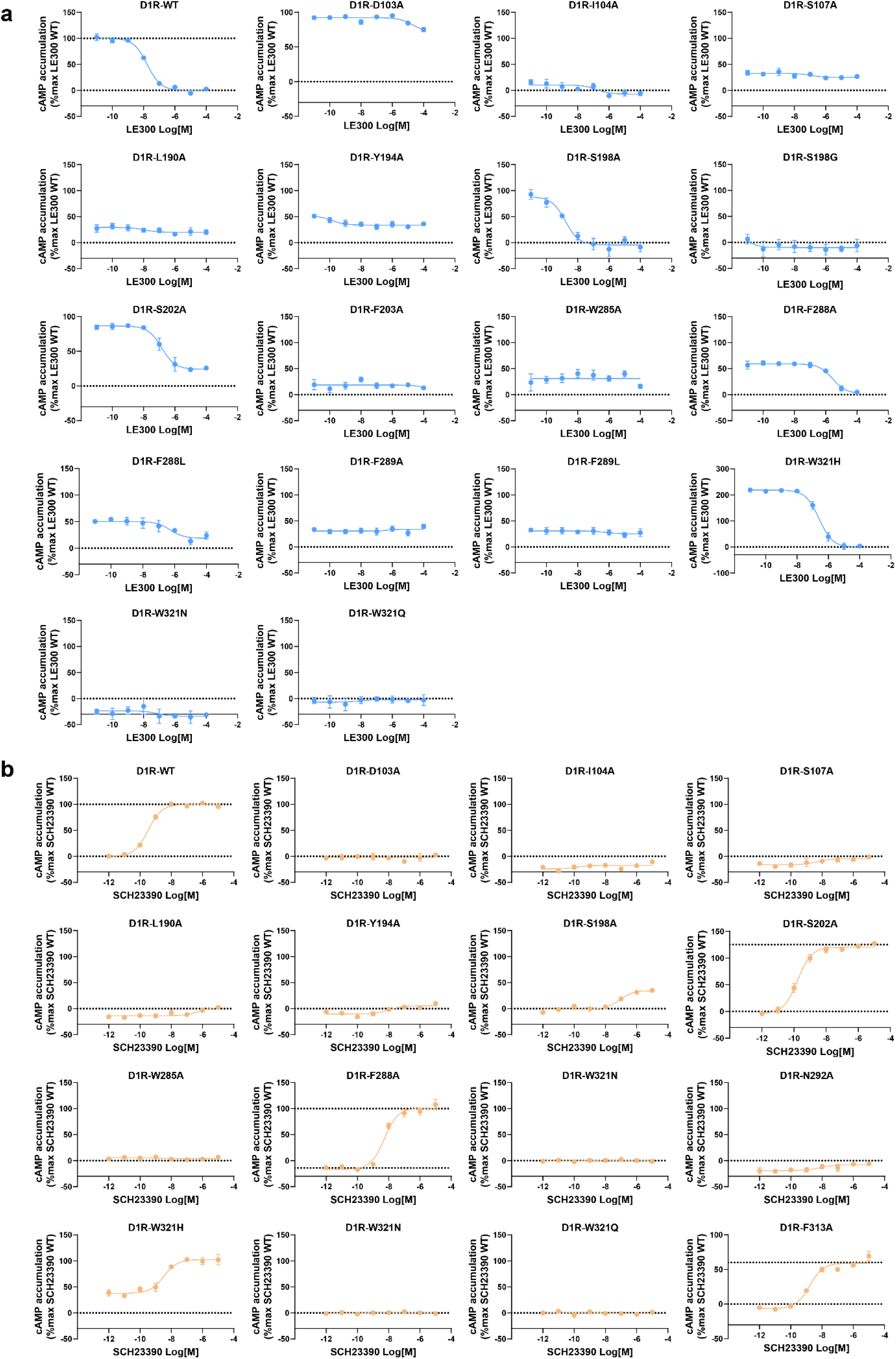
HTRF cAMP assay data of D1R wild-type and mutants. **a,** Concentration-response curves for dopamine and LE300 measured using the HTRF cAMP assay in HEK293 cells expressing the wild-type (WT) D1R. Responses were normalized to the maximal dopamine response (100%), with apo basal activity defined as 0%. Data are mean ± S.E.M. from three independent experiments (*n* = 3), each performed in triplicate. **b,** Concentration-response curves for dopamine and SCH23390 measured using the HTRF cAMP assay in HEK293 cells expressing the wild-type (WT) D1R. Responses were normalized to the maximal dopamine response (100%), with apo basal activity defined as 0%. Data are mean ± S.E.M. from three independent experiments (*n* = 3), each performed in triplicate.

**Extended Data Fig. 5.**
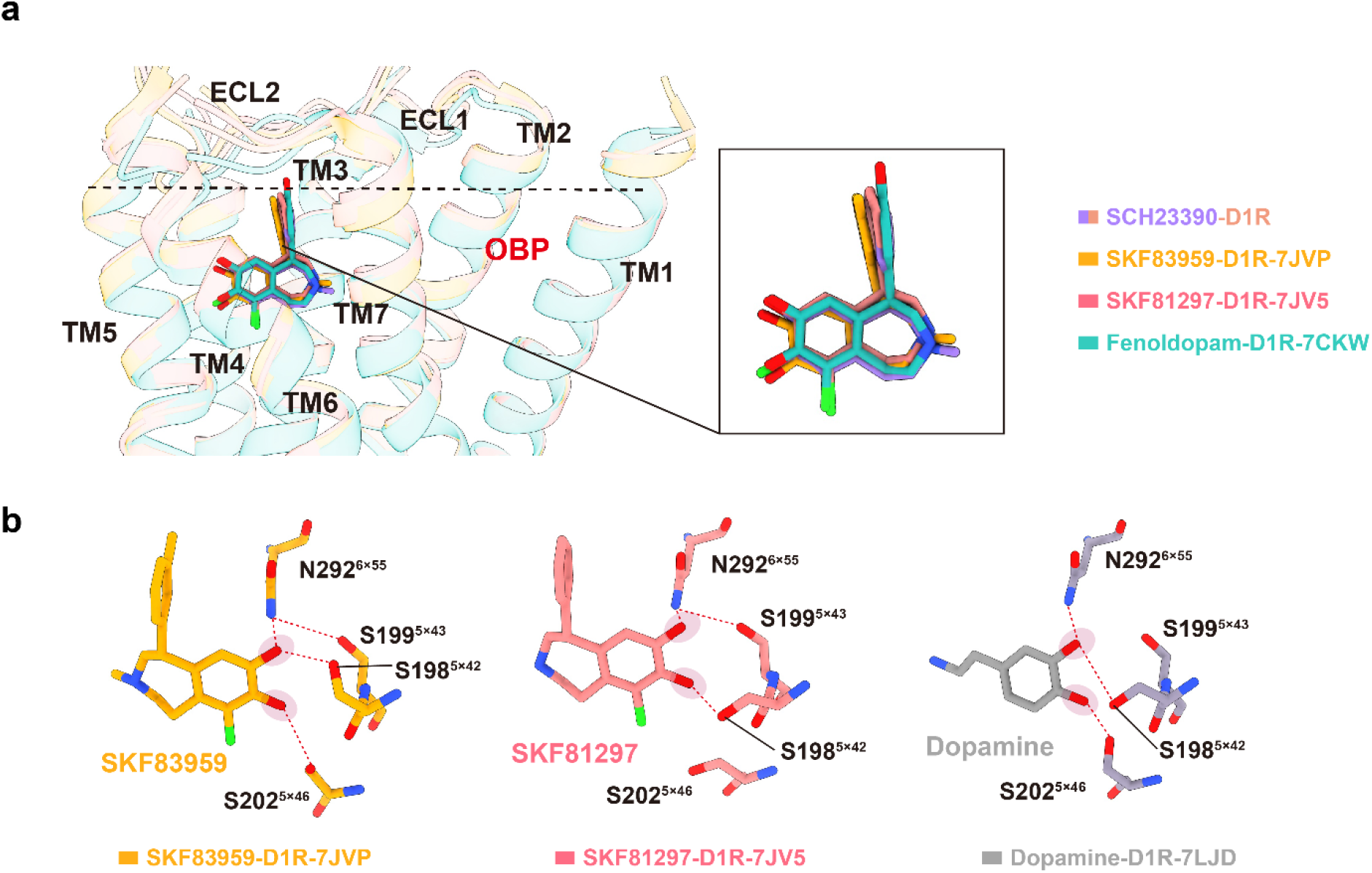
Conserved binding modes of benzazepine agonists in D1R. **a,** Superposition of SCH23390, SKF83959, SKF81297 and fenoldopam-bound D1R structures, showing a conserved L-shaped configuration of these benzazepine agonists within the orthosteric binding pocket. SKF83959, SKF81297 and fenoldopam-bound structures correspond to PDB 7JVP, 7JV5 and 7CKW, respectively. **b,** Comparison of the polar interaction networks formed by SKF83959, SKF81297 and dopamine in the catechol-binding region of D1R. Red dashed lines indicate polar interactions with N292^6.55^, S198^5.42^, S199^5.43^ and S202^5.46^. SKF83959, SKF81297 and dopamine-bound structures correspond to PDB 7JVP, 7JV5 and 7LJD, respectively.

**Extended Data Fig. 6.**
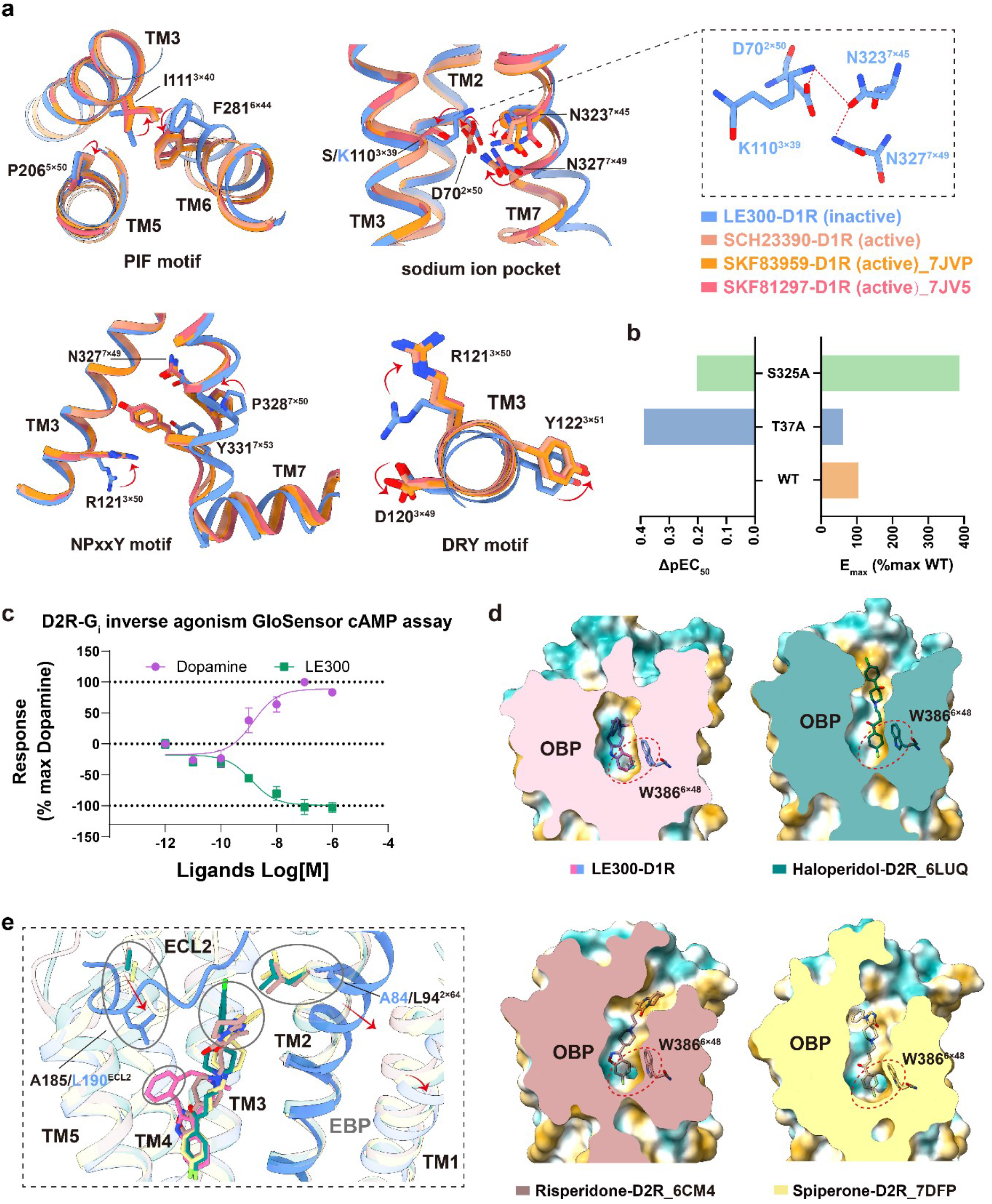
Structural basis of inverse agonism in LE300-bound D1R and inactive D2R complexes. **a,** P206^5.50^-I111^3.40^-F281^6.44^ (PIF), D120^3.49^-R121^3.50^-Y122^3.51^ (DRY), D70^2.50^-S110^3.39^- N323^7.45^-N327^7.49^ (sodium ion pocket) and N327^7.49^-P328^7.50^-xx-Y331^7.53^ (NPxxY) microswitches in LE300-bound inactive-state D1R and the three agonist-bound active-state complexes. Arrows denote conformational rearrangement. Notably, in the LE300-bound inactive-state D1R, the S110^3.39^K mutation enhances polar interactions with D70^2.50^ and N323^7.45^, thereby stabilizing the sodium ion pocket. **b,** Effects of key residues mutations T37^1×46^A and S325^7×47^A on SCH23390 induced cAMP responses in GloSensor cAMP assay. Responses were normalized to 100% for wild-type (ΔpEC_50_ = pEC_50_ of mutant − pEC_50_ of WT). **c,** Concentration-response curves for dopamine and LE300 measured using the GloSensor cAMP assay in HEK293 cells expressing WT D2R. Responses were normalized to the maximal dopamine response (100%), with apo basal activity defined as 0%. Data are mean ± S.E.M. from three independent experiments (*n* = 3), each performed in triplicate. **d,** The conserved toggle switch W285^6×48^ in the hydrophobic cavity of inverse agonist-binding pocket of D1R and D2R. Hydrophilic (blue) and hydrophobic (yellow) properties are shown. **e,** Side view superposition of LE300-bound inactive D1R with inactive D2R structures bound to haloperidol (PDB 6LUQ), risperidone (PDB 6CM4) or spiperone (PDB 7DFP). Circles highlight subtype-specific differences around A84^2.64^/L94^2.64^ and L190^ECL2^/A185^ECL2^ in D1R/D2R, respectively.

**Extended Data Fig. 7.**
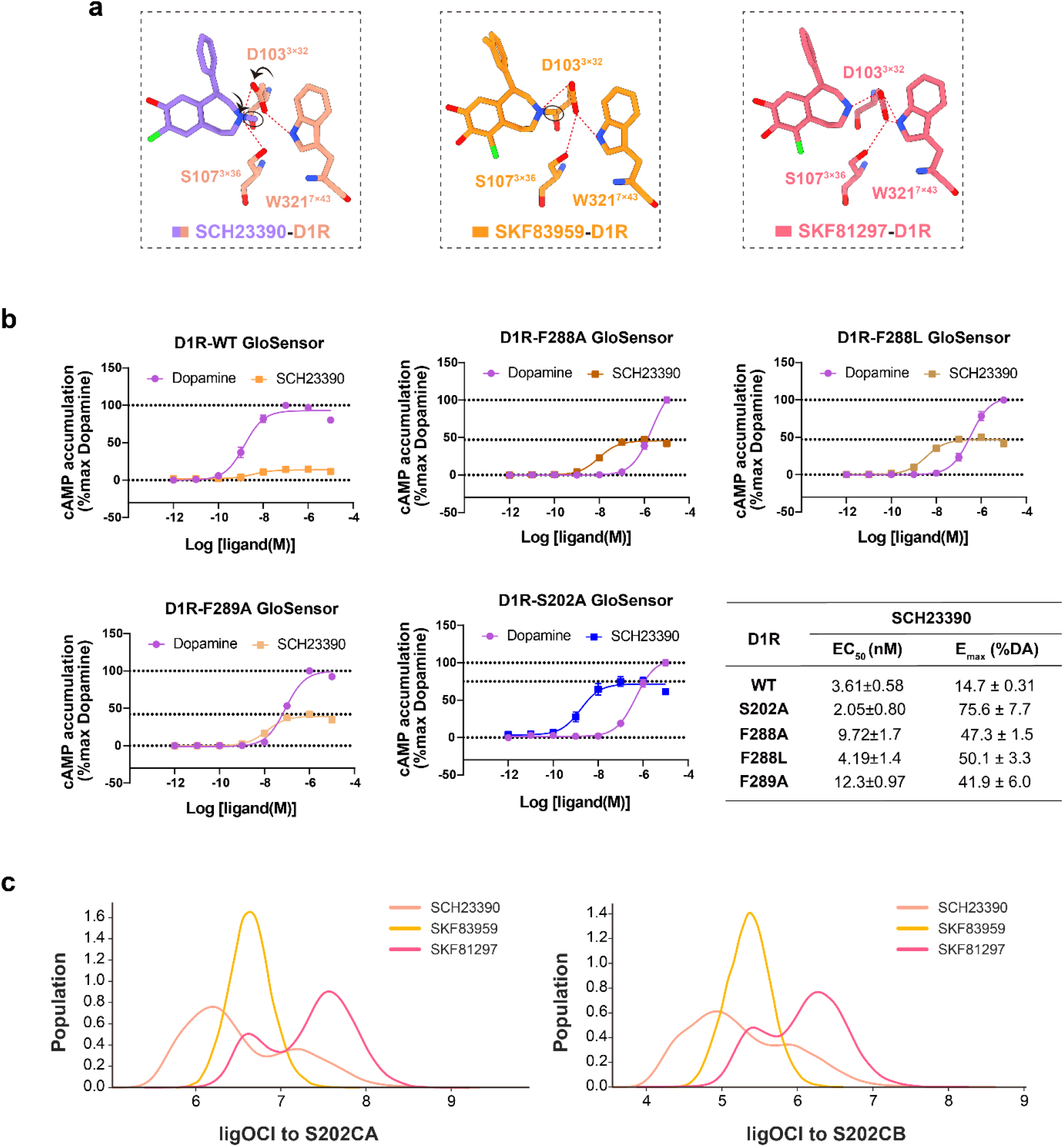
Orthosteric-pocket remodeling across graded D1R agonism. **a,** Structural comparison of the polar interaction network formed by D103^3.32^, S107^3.36^ and W321^7.43^ in SCH23390, SKF83959 and SKF81297-bound D1R. Red dashed lines indicate polar interactions, and arrows indicate relative side-chain (black) and helical (red) displacements. The arrow between the individual structures denotes increasing ligand efficacy. The SKF83959 and SKF81297-bound structures correspond to PDB 7JVP and 7JV5, respectively. **b,** Concentration-response curves for dopamine and SCH23390 at wild-type (WT) D1R and the S202^5.46^, F288^6.51^ and F289^6.52^ mutants, measured using the GloSensor cAMP assay. Responses were normalized to the maximal dopamine response. The table summarizes the potency (EC_50_) and maximal efficacy (Emax, expressed as a percentage of the maximal dopamine response) of SCH23390 at WT D1R and the mutants. Data are mean ± S.E.M. from three independent experiments (*n* = 3), each performed in triplicate. **c,** Distributions of the distances between the O/Cl atoms of SCH23390, SKF83959 or SKF81297 and the CA (left) or CB (right) atoms of S202^5.46^. These coordinates describe the position of the ligand relative to the extracellular portion of TM5. Distances are reported in Å.

**Extended Data Fig. 8.**
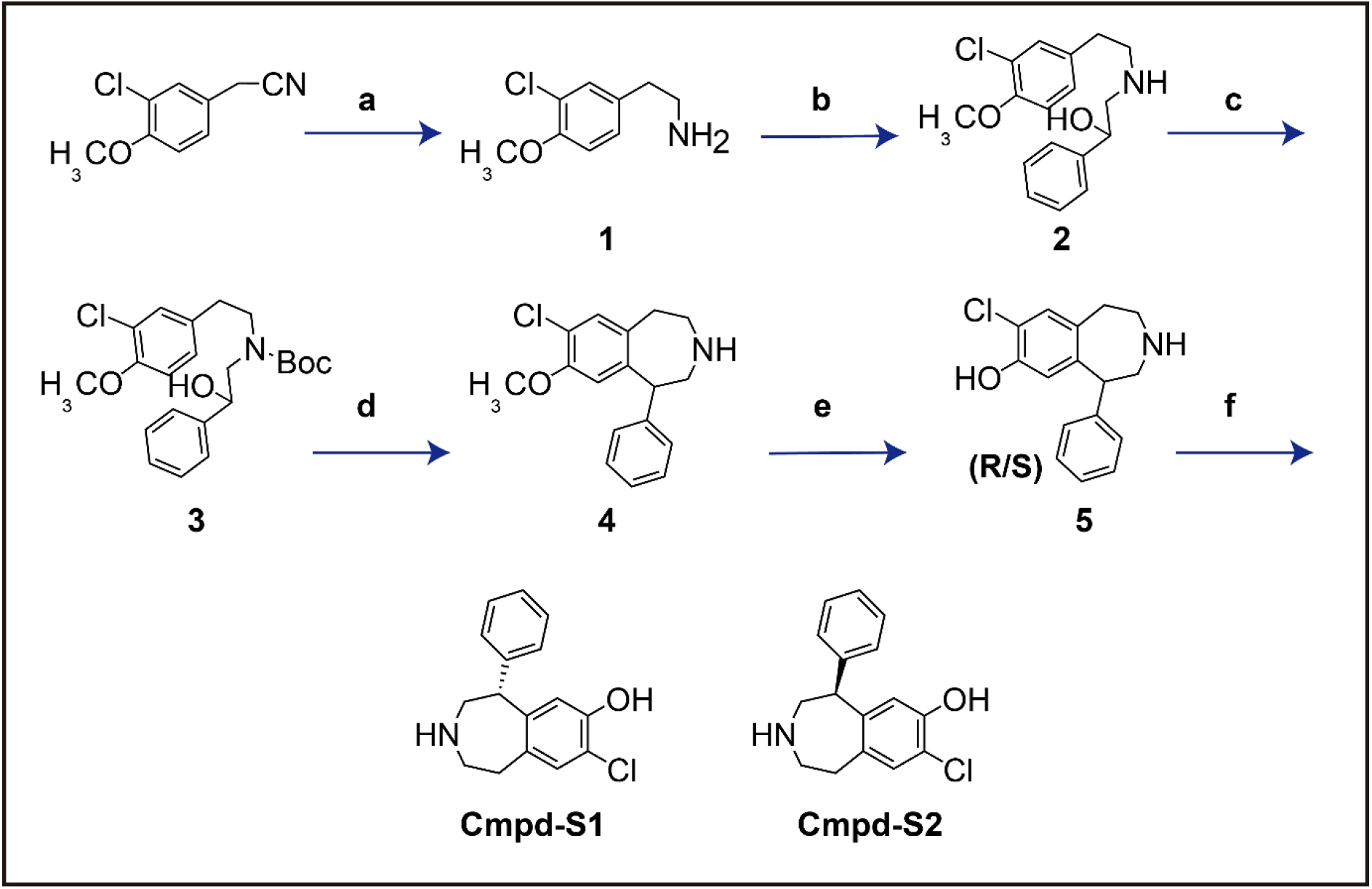
Synthesis of Cmpd-S1. Synthesis of compound 5 and its enantiomers Cmpd**-**S1 and Cmpd**-**S2. Reagents and conditions: (**a**) BH3-DMS, THF, reflux; (**b**) styreneoxide, AcOH, CH3CN, reflux; (**c**) Boc2O, Et3N, DCM; (**d**) H2SO4, TFA, reflux; (**e**)BBr3, DCM, −20 ℃; (**f**) chiral separation (AD-H column). Details shown in Methods.

**Supplementary Table 1.** Pharmacological efficacy spectrum of D1R and D2R ligands measured using the GloSensor cAMP assay (related to Figs. 1a, b and 5h, and Extended Data Fig. 6b).

| GloSensor cAMP assay |  |  |
| --- | --- | --- |
| Compounds | D1R-WT |  |
|  | EC<sub>50</sub> (nM) | E<sub>max</sub> (% Dopamine, DA) |
| Dopamine | 2.74±1.6 | 100 |
| SKF81297 | 2.09±1.6 | 106±2.8 |
| SKF83959 | 7.52±4.3 | 78.6±1.4 |
| SCH23390 | 4.85±0.95 | 26.4±6.9 |
| LE300 | 19.8±3.4 | -13.8±1.3 |
| Cmpd-S1 | 1.66±0.43 | 75.5±11 |
| Compounds | D2R-WT |  |
|  | EC<sub>50</sub> (nM) | E<sub>max</sub> (% Dopamine, DA) |
| Dopamine | 4.95±2.9 | 100 |
| LE300 | 1.19±0.13 | -103±6.8 |

Responses were normalized to the maximal dopamine response (100%), with apo basal activity defined as 0%. Data are mean ± S.E.M. from three independent experiments (*n* = 3), each performed in triplicate.

**Supplementary Table 2.**
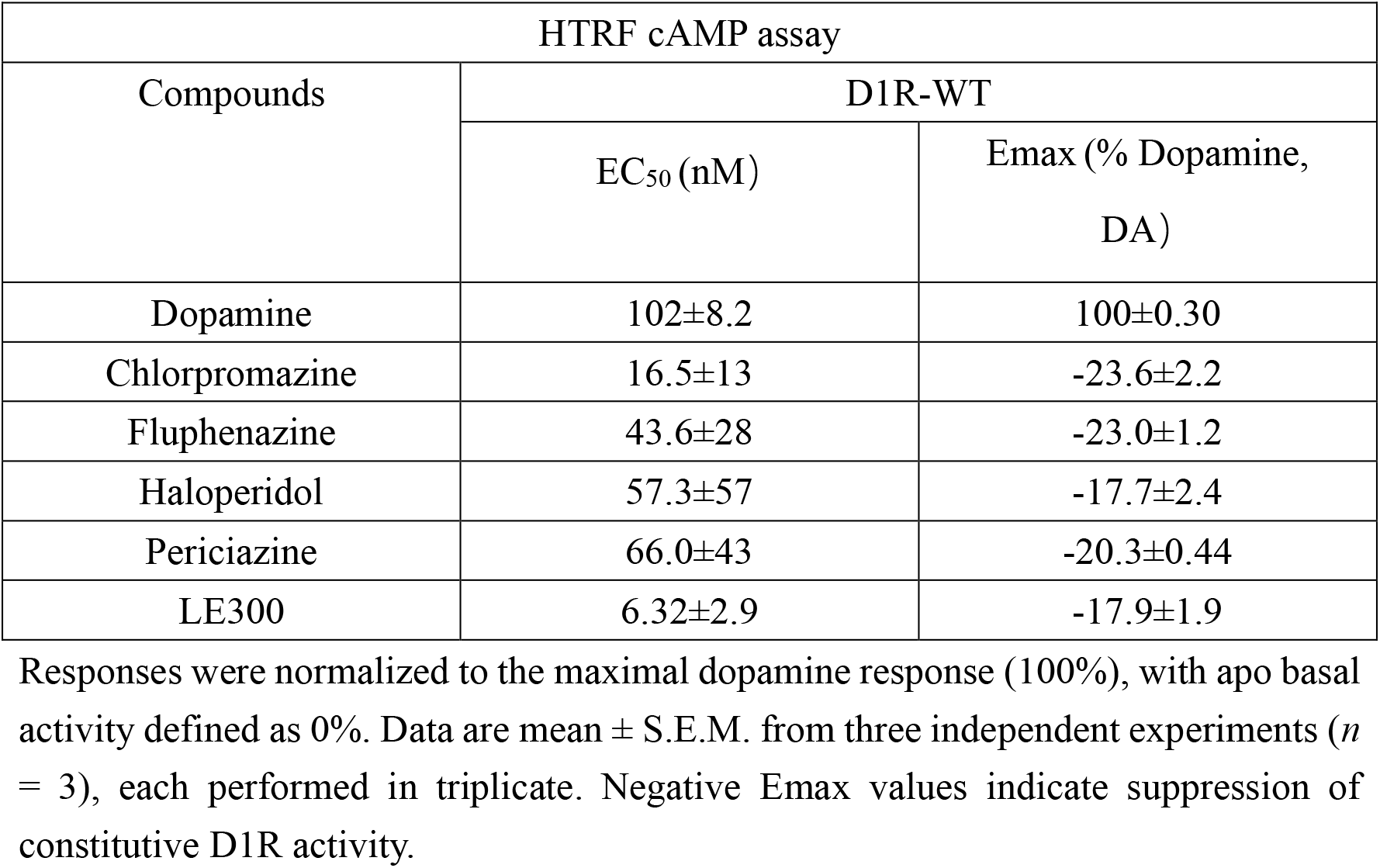
Effects of D1R inverse agonists measured using the HTRF cAMP assay (related to Extended Data Fig. 1).

**Supplementary Table 3.** Cryo-EM data collection, refinement, and validation (related to Figs. 2a, 3a and Extended Data Fig. 2).

|  | LE300-D1R | SCH23390-D1R-G<sub>s</sub>-Nb35 |
| --- | --- | --- |
| **Data collection and processing** |  |  |
| Magnification | 105,000 | 81,000 |
| Voltage (kV) | 300 | 300 |
| Electron exposure (e<sup>-</sup>/Å<sup>2</sup>) | 50 | 80 |
| Defocus range (μm) | -1.0~-2.0 | -1.0~-2.0 |
| Pixel size (Å) | 0.724 | 0.81 |
| Symmetry imposed | C1 | C1 |
| Initial particle projections (no.) | 3,835,046 | 2,734,371 |
| Final particle projections (no.) | 404,416 | 223,026 |
| Map resolution (Å) | 3.6 | 3.2 |
| Map resolution range (Å) | 2.8-6.0 | 2.4-5.0 |
| FSC threshold | 0.143 | 0.143 |
| **Model Refinement** |  |  |
| Real or reciprocal space | Real space | Real space |
| Model resolution (Å) | 3.6 | 3.2 |
| FSC threshold | 0.143 | 0.143 |
| B factors |  |  |
| Protein residues | 47.11/108.11/67.68 | 3.04/112.45/48.09 |
| Ligands | 57.15/57.15/57.15 | 41.52/69.55/59.49 |
| **Model composition** |  |  |
| Non-hydrogen atoms | 2070 | 8374 |
| Protein residues | 257 | 1041 |
| R.m.s. deviations |  |  |
| Bond lengths (Å) | 0.004 | 0.004 |
| Bond angles (°) | 0.932 | 0.950 |
| **Validation** |  |  |
| MolProbity score | 1.56 | 1.36 |
| Clashscore | 5.28 | 4.18 |
| Rotamer outliers (%) | 0.00 | 0.89 |
| Ramachandran plot |  |  |
| Favored (%) | 95.98 | 97.07 |
| Allowed (%) | 4.02 | 2.93 |
| Disallowed (%) | 0.00 | 0.00 |
| **Data availability** |  |  |
| EMDB entry | EMD-82510 | EMD-82511 |
| PDB entry | 44CW | 44CX |

**Supplementary Table 4.**
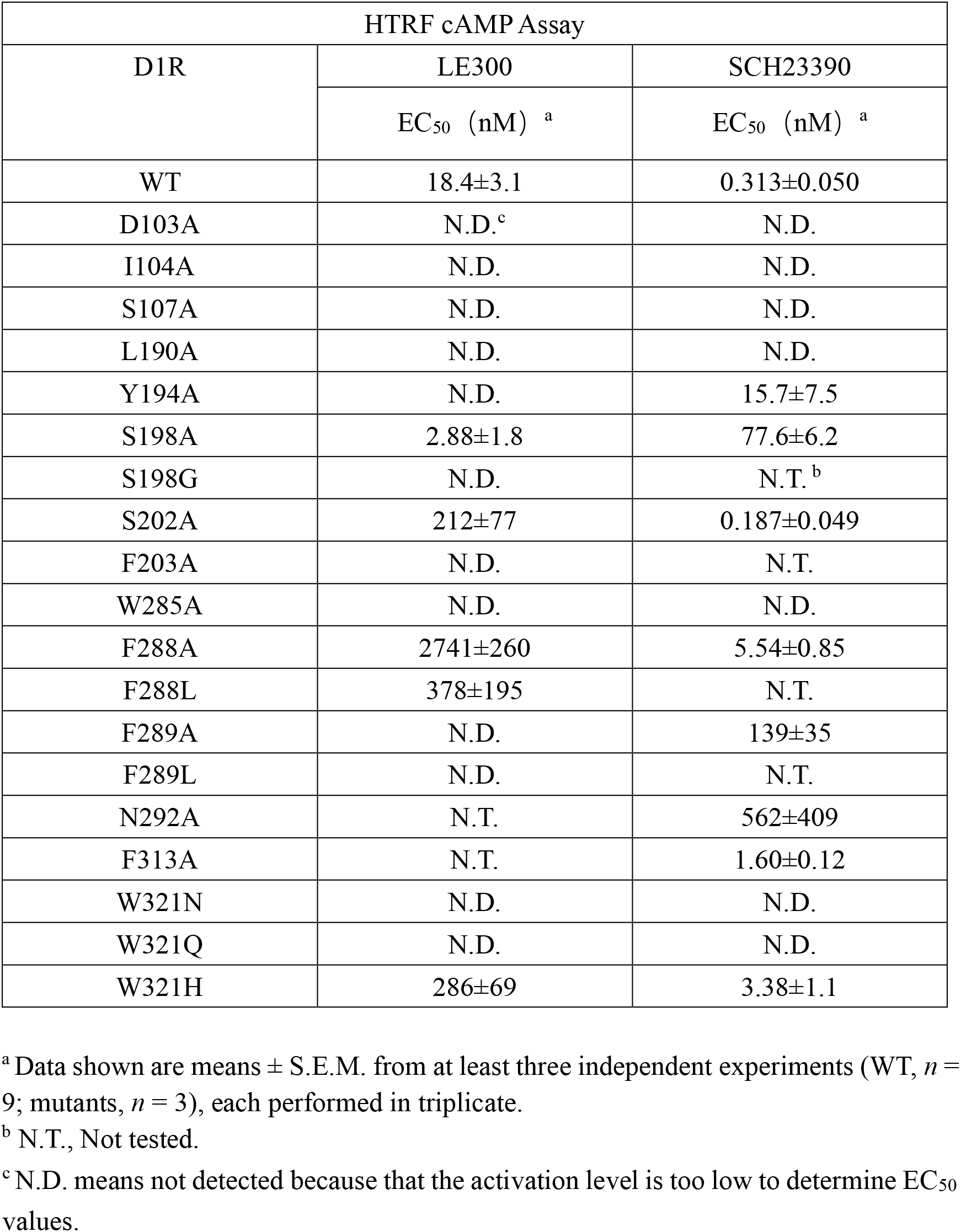
Potencies of LE300 and SCH23390 at wild-type and mutant D1R using the HTRF cAMP assay (related to Figs. 2d, 3b and Extended Data Fig. 4).

**Supplementary Table 5.**
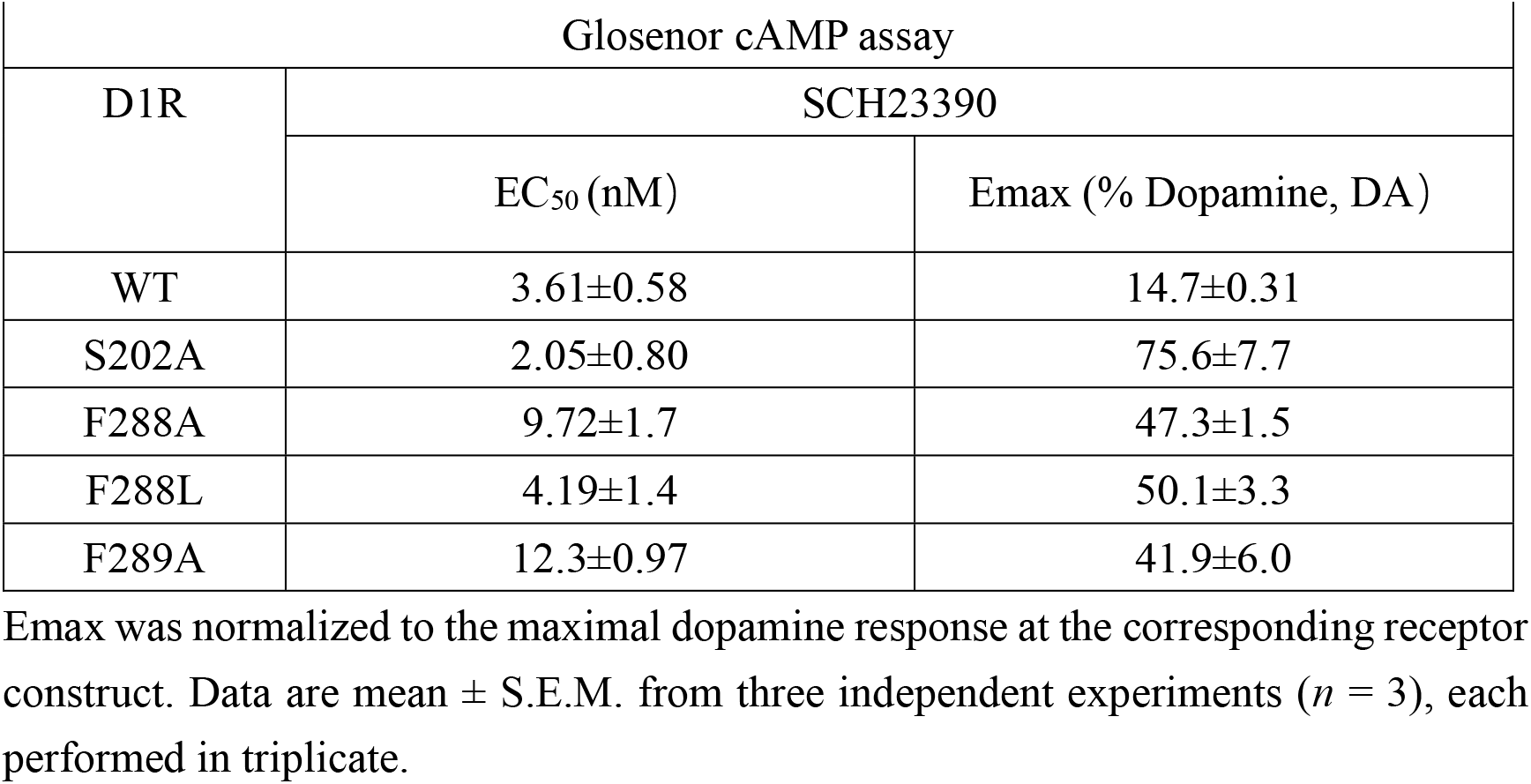
Effects of D1R mutations on SCH23390 potency and efficacy measured using the GloSensor cAMP assay (related to Fig. 5e and Extended Data Fig. 7c).

| GloSensor cAMP assay |  |  |
| --- | --- | --- |
| D1R | SCH23390 |  |
|  | EC<sub>50</sub> (nM) | E<sub>max</sub> (% Dopamine, DA) |
| WT | 3.61±0.58 | 14.7±0.31 |
| S202A | 2.05±0.80 | 75.6±7.7 |
| F288A | 9.72±1.7 | 47.3±1.5 |
| F288L | 4.19±1.4 | 50.1±3.3 |
| F289A | 12.3±0.97 | 41.9±6.0 |

E<sub>max</sub> was normalized to the maximal dopamine response at the corresponding receptor construct. Data are mean ± S.E.M. from three independent experiments (*n* = 3), each performed in triplicate.

**Supplementary Table 6.**
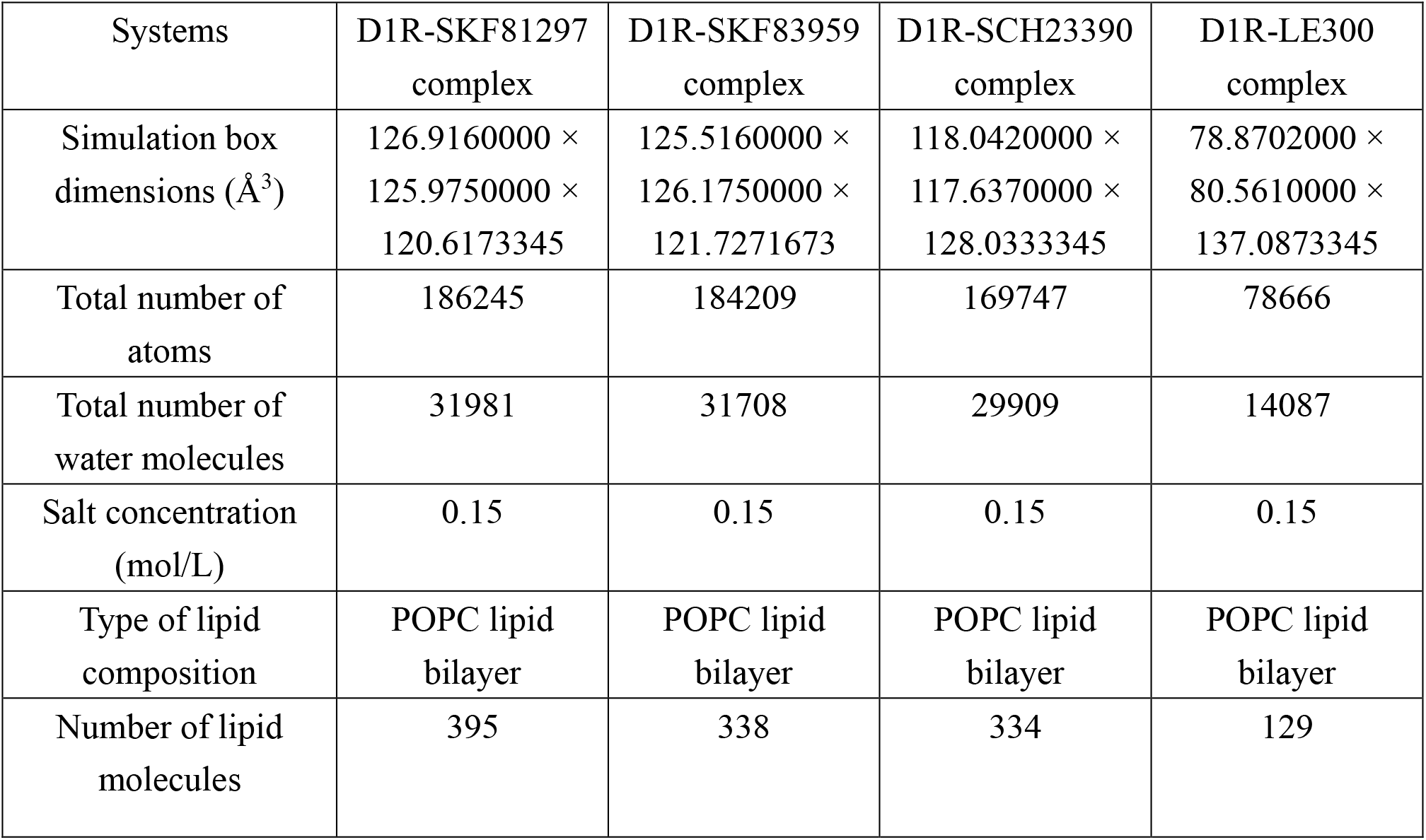
System setup for molecular dynamics simulations (related to Fig. 5g and Extended Data Fig. 7c).

| Systems | D1R-SKF81297<br>complex | D1R-SKF83959<br>complex | D1R-SCH23390<br>complex | D1R-LE300<br>complex |
| --- | --- | --- | --- | --- |
| Simulation box<br>dimensions ($\text{\AA}^3$) | 126.9160000 ×<br>125.9750000 ×<br>120.6173345 | 125.5160000 ×<br>126.1750000 ×<br>121.7271673 | 118.0420000 ×<br>117.6370000 ×<br>128.0333345 | 78.8702000 ×<br>80.5610000 ×<br>137.0873345 |
| Total number of<br>atoms | 186245 | 184209 | 169747 | 78666 |
| Total number of<br>water molecules | 31981 | 31708 | 29909 | 14087 |
| Salt concentration<br>(mol/L) | 0.15 | 0.15 | 0.15 | 0.15 |
| Type of lipid<br>composition | POPC lipid<br>bilayer | POPC lipid<br>bilayer | POPC lipid<br>bilayer | POPC lipid<br>bilayer |
| Number of lipid<br>molecules | 395 | 338 | 334 | 129 |

## References

1 Beaulieu, J. M. & Gainetdinov, R. R. The physiology, signaling, and pharmacology of dopamine receptors. Pharmacological reviews 63, 182–217 (2011). 10.1124/pr.110.002642

2 Missale, C., Nash, S. R., Robinson, S. W., Jaber, M. & Caron, M. G. Dopamine receptors: From structure to function. Physiological reviews 78, 189–225 (1998). 10.1152/physrev.1998.78.1.189

3 Klein, M. O. et al. Dopamine: Functions, Signaling, and Association with Neurological Diseases. Cell Mol Neurobiol 39, 31–59 (2019). 10.1007/s10571-018-0632-3

4 Zhuang, Y. et al. Structural insights into the human D1 and D2 dopamine receptor signaling complexes. Cell 184, 931–942.e918 (2021). 10.1016/j.cell.2021.01.027

5 Wood, M., Dubois, V., Scheller, D. & Gillard, M. Rotigotine is a potent agonist at dopamine D1 receptors as well as at dopamine D2 and D3 receptors. British Journal of Pharmacology 172, 1124–1135 (2015). 10.1111/bph.12988

6 Yegla, B., Rubin, J., Grall, M., Formella, A. & Jenner, P. Apomorphine differentially engages the cAMP and beta-arrestin signaling pathways relative to dopamine at human dopamine receptors. J Neural Transm (Vienna) 132, 1751–1760 (2025). 10.1007/s00702-025-03042-7

7 Cai, G., Gurdal, H., Smith, C., Wang, H. Y. & Friedman, E. Inverse agonist properties of dopaminergic antagonists at the D(1A) dopamine receptor: uncoupling of the D(1A) dopamine receptor from G(s) protein. Mol Pharmacol 56, 989–996 (1999). 10.1124/mol.56.5.989

8 Cai, G., Gurdal, H., Smith, C., Wang, H.-Y. & Friedman, E. Inverse Agonist Properties of Dopaminergic Antagonists at the D1A Dopamine Receptor: Uncoupling of the D1ADopamine Receptor from Gs Protein. Molecular Pharmacology 56, 989–996 (1999). 10.1016/S0026-895X(24)12778-2

9 Bourne, J. A. SCH 23390: The First Selective Dopamine D1-Like Receptor Antagonist. CNS Drug Reviews 7, 399–414 (2001). 10.1111/j.1527-3458.2001.tb00207.x

10 Waddington, J. L. & O’Boyle, K. M. Drugs acting on brain dopamine receptors: A conceptual re-evaluation five years after the first selective D-1 antagonist. Pharmacology & Therapeutics 43, 1–52 (1989). 10.1016/0163-7258(89)90046-6

11 O’Boyle, K. M. & Waddington, J. L. Agonist and antagonist interactions with D1 dopamine receptors: agonist-induced masking of D1 receptors depends on intrinsic activity. Neuropharmacology 31, 177–183 (1992). 10.1016/0028-3908(92)90029-o

12 Kassack, M. U., Höfgen, B., Decker, M., Eckstein, N. & Lehmann, J. Pharmacological characterization of the benz[d]indolo[2,3-g]azecine LE300, a novel type of a nanomolar dopamine receptor antagonist. Naunyn-Schmiedeberg’s Archives of Pharmacology 366, 543–550 (2002). 10.1007/s00210-002-0641-z

13 Lee, S.-M. et al. SKF-83959 is not a highly-biased functionally selective D1 dopamine receptor ligand with activity at phospholipase C. Neuropharmacology 86, 145–154 (2014). 10.1016/j.neuropharm.2014.05.042

14 Mannoury la Cour, C., Vidal, S., Pasteau, V., Cussac, D. & Millan, M. J. Dopamine D1 receptor coupling to Gs/olf and Gq in rat striatum and cortex: A scintillation proximity assay (SPA)/antibody-capture characterization of benzazepine agonists. Neuropharmacology 52, 1003–1014 (2007). 10.1016/j.neuropharm.2006.10.021

15 Nguyen, A. M. et al. Characterization of Gαs and Gαolf activation by catechol and non-catechol dopamine D1 receptor agonists. iScience 28, 112345 (2025). 10.1016/j.isci.2025.112345

16 Bezard, E. et al. Rationale and Development of Tavapadon, a D1/D5-Selective Partial Dopamine Agonist for the Treatment of Parkinson’s Disease. CNS Neurol Disord Drug Targets 23, 476–487 (2024). 10.2174/1871527322666230331121028

17 Lewis, M. M., Watts, V. J., Lawler, C. P., Nichols, D. E. & Mailman, R. B. Homologous desensitization of the D1A dopamine receptor: efficacy in causing desensitization dissociates from both receptor occupancy and functional potency. J Pharmacol Exp Ther 286, 345–353 (1998).

18 Wade, M. R. & Nomikos, G. G. Tolerance to the procholinergic action of the D1 receptor full agonist dihydrexidine. Psychopharmacology (Berl) 182, 393–399 (2005). 10.1007/s00213-005-0106-4

19 Riesenberg, R., Werth, J., Zhang, Y., Duvvuri, S. & Gray, D. PF-06649751 efficacy and safety in early Parkinson’s disease: a randomized, placebo-controlled trial. Ther Adv Neurol Disord 13, 1756286420911296 (2020). 10.1177/1756286420911296

20 Sohur, U. S. et al. Phase 1 Parkinson’s Disease Studies Show the Dopamine D1/D5 Agonist PF-06649751 is Safe and Well Tolerated. Neurol Ther 7, 307–319 (2018). 10.1007/s40120-018-0114-z

21 Zhuang, Y. et al. Mechanism of dopamine binding and allosteric modulation of the human D1 dopamine receptor. Cell Research 31, 593–596 (2021). 10.1038/s41422-021-00482-0

22 Teng, X. et al. Ligand recognition and biased agonism of the D1 dopamine receptor. Nature Communications 13, 3186 (2022). 10.1038/s41467-022-30929-w

23 Xu, P. et al. Structural genomics of the human dopamine receptor system. Cell Research 33, 604–616 (2023). 10.1038/s41422-023-00808-0

24 Sibley, D. R., Luderman, K. D., Free, R. B. & Shi, L. Novel Cryo-EM structures of the D1 dopamine receptor unlock its therapeutic potential. Signal Transduction and Targeted Therapy 6, 205 (2021). 10.1038/s41392-021-00630-3

25 Xiao, P. et al. Ligand recognition and allosteric regulation of DRD1-Gs signaling complexes. Cell 184, 943–956.e918 (2021). 10.1016/j.cell.2021.01.028

26 Teng, X. et al. Structural insights into G protein activation by D1 dopamine receptor. Science Advances 8, eabo4158 10.1126/sciadv.abo4158

27 Mahan, L. C., Burch, R. M., Monsma, F. J., Jr. & Sibley, D. R. Expression of striatal D1 dopamine receptors coupled to inositol phosphate production and Ca2+ mobilization in Xenopus oocytes. Proceedings of the National Academy of Sciences of the United States of America 87, 2196–2200 (1990). 10.1073/pnas.87.6.2196

28 Zhuang, Y. et al. Structural insights into the human D1 and D2 dopamine receptor signaling complexes. Cell 184, 931–942 e918 (2021). 10.1016/j.cell.2021.01.027

29 Fan, L. et al. Haloperidol bound D2 dopamine receptor structure inspired the discovery of subtype selective ligands. Nature Communications 11, 1074 (2020). 10.1038/s41467-020-14884-y

30 Wang, S. et al. Structure of the D2 dopamine receptor bound to the atypical antipsychotic drug risperidone. Nature 555, 269–273 (2018). 10.1038/nature25758

31 Im, D. et al. Structure of the dopamine D2 receptor in complex with the antipsychotic drug spiperone. Nature Communications 11, 6442 (2020). 10.1038/s41467-020-20221-0

32 Nygaard, R. et al. The dynamic process of beta(2)-adrenergic receptor activation. Cell 152, 532–542 (2013). 10.1016/j.cell.2013.01.008

33 Manglik, A. et al. Structural Insights into the Dynamic Process of beta2-Adrenergic Receptor Signaling. Cell 161, 1101–1111 (2015). 10.1016/j.cell.2015.04.043

34 Eddy, M. T. et al. Allosteric Coupling of Drug Binding and Intracellular Signaling in the A(2A) Adenosine Receptor. Cell 172, 68–80 e12 (2018). 10.1016/j.cell.2017.12.004

35 Weis, W. I. & Kobilka, B. K. The Molecular Basis of G Protein-Coupled Receptor Activation. Annu Rev Biochem 87, 897–919 (2018). 10.1146/annurev-biochem-060614-033910

36 Su, M. et al. Structures of beta(1)-adrenergic receptor in complex with Gs and ligands of different efficacies. Nature communications 13, 4095 (2022). 10.1038/s41467-022-31823-1

37 Yang, F. et al. Different conformational responses of the β2-adrenergic receptor-Gs complex upon binding of the partial agonist salbutamol or the full agonist isoprenaline. National science review 8 (2021). 10.1093/nsr/nwaa284

38 Liu, P. et al. The structural basis of the dominant negative phenotype of the Gαi1β1γ2 G203A/A326S heterotrimer. Acta Pharmacologica Sinica 37, 1259–1272 (2016). 10.1038/aps.2016.69

39 Carpenter, B., Nehmé, R., Warne, T., Leslie, A. G. W. & Tate, C. G. Structure of the adenosine A2A receptor bound to an engineered G protein. Nature 536, 104–107 (2016). 10.1038/nature18966

40 García-Nafría, J., Lee, Y., Bai, X., Carpenter, B. & Tate, C. G. Cryo-EM structure of the adenosine A(2A) receptor coupled to an engineered heterotrimeric G protein. Elife 7 (2018). 10.7554/eLife.35946

41 Chen, H., Huang, W. & Li, X. Structures of oxysterol sensor EBI2/GPR183, a key regulator of the immune response. Structure 30, 1016–1024.e1015 (2022). 10.1016/j.str.2022.04.006

42 Rasmussen, S. G. et al. Crystal structure of the β2 adrenergic receptor-Gs protein complex. Nature 477, 549–555 (2011). 10.1038/nature10361

43 Maeda, S. et al. Development of an antibody fragment that stabilizes GPCR/G-protein complexes. Nature Communications 9, 3712 (2018). 10.1038/s41467-018-06002-w

44 Mastronarde, D. N. Automated electron microscope tomography using robust prediction of specimen movements. J Struct Biol 152, 36–51 (2005). 10.1016/j.jsb.2005.07.007

45 Zheng, S. Q. et al. MotionCor2: anisotropic correction of beam-induced motion for improved cryo-electron microscopy. Nature methods 14, 331–332 (2017). 10.1038/nmeth.4193

46 Punjani, A., Rubinstein, J. L., Fleet, D. J. & Brubaker, M. A. cryoSPARC: algorithms for rapid unsupervised cryo-EM structure determination. Nature Methods 14, 290–296 (2017). 10.1038/nmeth.4169

47 Zivanov, J. et al. New tools for automated high-resolution cryo-EM structure determination in RELION-3. eLife 7, e42166 (2018). 10.7554/eLife.42166

48 Zhang, K. Gctf: Real-time CTF determination and correction. Journal of Structural Biology 193, 1–12 (2016). 10.1016/j.jsb.2015.11.003

49 Emsley, P. & Cowtan, K. Coot: model-building tools for molecular graphics. Acta crystallographica. Section D, Biological crystallography 60, 2126–2132 (2004). 10.1107/S0907444904019158

50 Schott-Verdugo, S. & Gohlke, H. PACKMOL-Memgen: A Simple-To-Use, Generalized Workflow for Membrane-Protein–Lipid-Bilayer System Building. Journal of Chemical Information and Modeling 59, 2522–2528 (2019). 10.1021/acs.jcim.9b00269

51 Tian, C. et al. ff19SB: Amino-Acid-Specific Protein Backbone Parameters Trained against Quantum Mechanics Energy Surfaces in Solution. Journal of chemical theory and computation 16, 528–552 (2020). 10.1021/acs.jctc.9b00591

52 Dickson, C. J., Walker, R. C. & Gould, I. R. Lipid21: Complex Lipid Membrane Simulations with AMBER. Journal of Chemical Theory and Computation 18, 1726–1736 (2022). 10.1021/acs.jctc.1c01217

53 He, X., Man, V. H., Yang, W., Lee, T. S. & Wang, J. A fast and high-quality charge model for the next generation general AMBER force field. J Chem Phys 153, 114502 (2020). 10.1063/5.0019056

54 Lu, S. et al. Activation pathway of a G protein-coupled receptor uncovers conformational intermediates as targets for allosteric drug design. Nat Commun 12, 4721 (2021). 10.1038/s41467-021-25020-9

55 He, X. et al. Conformational Selection Mechanism Provides Structural Insights into the Optimization of APC-Asef Inhibitors. Molecules 26 (2021). 10.3390/molecules26040962

56 Salomon-Ferrer, R., Gotz, A. W., Poole, D., Le Grand, S. & Walker, R. C. Routine Microsecond Molecular Dynamics Simulations with AMBER on GPUs. 2. Explicit Solvent Particle Mesh Ewald. Journal of chemical theory and computation 9, 3878–3888 (2013). 10.1021/ct400314y

